# A Next Generation Connectivity Map: L1000 Platform And The First 1,000,000 Profiles

**DOI:** 10.1101/136168

**Authors:** Aravind Subramanian, Rajiv Narayan, Steven M. Corsello, David D. Peck, Ted E. Natoli, Xiaodong Lu, Joshua Gould, John F. Davis, Andrew A. Tubelli, Jacob K. Asiedu, David L. Lahr, Jodi E. Hirschman, Zihan Liu, Melanie Donahue, Bina Julian, Mariya Khan, David Wadden, Ian Smith, Daniel Lam, Arthur Liberzon, Courtney Toder, Mukta Bagul, Marek Orzechowski, Oana M. Enache, Federica Piccioni, Alice H. Berger, Alykhan Shamji, Angela N. Brooks, Anita Vrcic, Corey Flynn, Jacqueline Rosains, David Takeda, Desiree Davison, Justin Lamb, Kristin Ardlie, Larson Hogstrom, Nathanael S. Gray, Paul A. Clemons, Serena Silver, Xiaoyun Wu, Wen-Ning Zhao, Willis Read-Button, Xiaohua Wu, Stephen J. Haggarty, Lucienne V. Ronco, Jesse S. Boehm, Stuart L. Schreiber, John G. Doench, Joshua A. Bittker, David E. Root, Bang Wong, Todd R. Golub

## Abstract

We previously piloted the concept of a Connectivity Map (CMap), whereby genes, drugs and disease states are connected by virtue of common gene-expression signatures. Here, we report more than a 1,000-fold scale-up of the CMap as part of the NIH LINCS Consortium, made possible by a new, low-cost, high throughput reduced representation expression profiling method that we term L1000. We show that L1000 is highly reproducible, comparable to RNA sequencing, and suitable for computational inference of the expression levels of 81% of non-measured transcripts. We further show that the expanded CMap can be used to discover mechanism of action of small molecules, functionally annotate genetic variants of disease genes, and inform clinical trials. The 1.3 million L1000 profiles described here, as well as tools for their analysis, are available at https://clue.io.

**HIGHLIGHTS:**

- A new gene expression profiling method, L1000, dramatically lowers cost
- The Connectivity Map database now includes 1.3 million publicly accessible L1000 perturbational profiles
- This expanded Connectivity Map facilitates discovery of small molecule mechanism of action and functional annotation of genetic variants
- The work establishes feasibility and utility of a truly comprehensive Connectivity Map

## INTRODUCTION

The sequencing of the human genome provided the parts list of life, and this in turn has led to an explosion of new insights into the genetic basis of disease. Genome-wide association studies have identified risk-associated loci for major diseases, and the sequencing of human tumors has similarly identified the somatic mutations that underlie many types of cancer. The research community has benefitted from these genomic resources by being able to readily look up sequence variants in large-scale compendia of genomic variation. Such look-up tables of biology, which support the generation and interrogation of new hypotheses, have transformed how modern research is done.

A challenge, however, is that a parts list and its association with disease is generally not sufficient to establish causality, and to provide mechanistic and circuit-level insights into uncharacterized gene products. Even beyond the precise biochemical function of particular proteins, the pathways in which they operate are often unknown. Truly understanding cellular function requires perturbing the system – modulating the expression of a gene of interest, and monitoring the downstream consequences. Of course the detailed characterization of individual proteins – one at a time – is the bedrock of biomedical research. But systematic, large-scale compendia of the cellular effects of genetic perturbation have yet to be established as a community resource.

Similarly, there has been no method to systematically determine the cellular effects of a given chemical compound. For example, it would be desirable to be able to query a functional look-up table to discover unexpected off-target activities of a compound – such activities often being discovered only late in the drug development process, resulting in side effects that limit clinical use. Furthermore, the mechanism of action of drugs with proven clinical benefit is often poorly understood, thereby making it difficult to develop nextgeneration drugs that improve their utility.

We previously hypothesized that a potential solution to these problems might be the creation of a comprehensive catalog of cellular signatures representing systematic perturbation with genetic perturbagens (reflecting protein function) and pharmacologic perturbagens (reflecting small molecule function). Signatures that proved to be similar might thus represent useful and previously unrecognized connections (e.g., between two proteins operating in the same pathway, between a small molecule and its protein target, or between two small molecules of similar function but structural dissimilarity). Such a catalog of connections could thus serve as a functional look-up table of the genome, and we termed this concept the Connectivity Map (Lamb, 2006).

We previously piloted the Connectivity Map (CMap) concept by treating cells with 164 FDA-approved drugs and tool compounds, and then performing mRNA expression profiling using Affymetrix microarrays. A total of 564 gene expression profiles were generated, and a measure of gene expression signature similarity based on the Kolmogorov-Smirnov statistic was developed so that users could query the CMap database using a userdefined signature of interest (e.g., a signature of disease state, cellular process, genetic perturbation or small molecule action). The pilot Connectivity Map database has served as a public resource, with over 18,000 registered users who have submitted over 150,000 queries to the CMap website (broadinstitute.org/cmap).

Recently published uses of the CMap pilot dataset include discovery of the anthelmintic drug parbendazole as an inducer of osteoclast differentiation (Brum et al., 2015), the triterpene celastrol as a leptin sensitizer (Liu et al., 2015), compounds targeting COX2 and ADRA2A as potential diabetes treatments (Zhang et al., 2015), small molecule therapeutics for skeletal muscular atrophy (Dyle et al., 2014) and spinal muscular atrophy (Farooq et al., 2009), and new therapeutic hypotheses for the treatment of inflammatory bowel disease (Dudley et al., 2011) and cancer (Singh et al., 2016); (Muthuswami et al., 2013; Wang et al., 2008); (Schnell et al., 2015); (Fortney et al., 2015; Wang et al., 2011); (Vilar et al., 2009); (Churchman et al., 2015).

Recent reports similarly point to the utility of CMap in dissecting biological pathways, including the identification of regulators of the transcription factor p73 (Rosenbluth et al., 2008), the discovery of pharmacologic modulators of the unfolded-protein response (Saito et al., 2009), and the discovery of the mechanism of action of the cytostatic compound CCT020312 as a regulator a translation initiation factor kinase known as EIF2AK3 (Stockwell et al., 2012).

Despite the popularity of the pilot Connectivity Map pilot dataset, its small scale limits its utility. With only 164 drug perturbations in only 3 cancer cell lines, the database lacks the necessary richness of a truly genome-scale resource. Missing is a diversity of chemical perturbations, genetic perturbations (i.e., gain-and loss-of-function studies) as well as a diversity of cell types that ideally would span multiple cell lineages and cell states. Ultimately, a Connectivity Map should contain signatures of perturbation of all genes in the genome, all small molecules of interest to the community, and all cell types of relevance to biomedical research. Such a database might need to contain tens of millions of gene expression profiles to be truly comprehensive. Unfortunately, the high cost of commercial gene expression microarrays and even RNA sequencing precludes such a genome-scale Connectivity Map. We therefore describe here a new approach to gene expression profiling based on a reduced representation of the transcriptome. This method, which we call L1000, is high-throughput and low-cost, and is thus well-suited to a large-scale Connectivity Map. We also report here the first 1,319,138 L1000 profiles as part of the NIH LINCS initiative, representing over a 1,000-fold expansion of the pilot Connectivity Map resource.

## RESULTS

### Reduced representation of transcriptome

Some of the earliest studies of genome-wide expression data showed that gene expression is highly correlated, with clusters of genes exhibiting similar expression patterns across cell states (Eisen et al., 1998). Given such correlation structure, we hypothesized that it might be possible to capture at low cost any cellular state by measuring a reduced representation of the transcriptome. This concept is similar to that used routinely in human genetics, whereby a reduced representation of genetic variation is used to guide genotyping of a ‘tag SNP’ within a haplotype block and then the other (non-measured) SNPs on the haplotype are computationally inferred.

To explore this transcriptional reduced representation concept, we first assembled, from NCBI’s Gene Expression Omnibus (GEO) (Edgar et al., 2002), a diverse collection of 12,031 gene expression profiles generated using Affymetrix HGU133A arrays. We used these to identify a subset of informative transcripts, which we term ‘landmark’ transcripts. We sought to determine the optimal number of landmarks *k*. If *k* was too small, too much information might be lost, whereas if *k* was too large, sufficient cost reduction compared to the entire transcriptome might be not be achieved. To address this, we asked what number of landmarks would optimally recover the observed connections seen in the pilot Connectivity Map dataset based on Affymetrix arrays (Dataset DS_CMAP-AFFX_). Specifically, our prior work indicated that 25 query signatures yielded robust and expected connections to small molecules in the CMap pilot dataset (Table S1). We therefore used those 25 signatures to query the imputed DS_CMAP-AFFX_ dataset for each value of *k*, counting how often we recovered the connections observed in the original dataset at a comparable rank based on the Kolmogorov-Smirnov statistic (See Methods). Figure S1A shows the results of this analysis, which revealed that 1,000 landmarks were sufficient to recover 82% of expected connections. This computational simulation provided an early indication that measuring a subset of the transcriptome might recapitulate much of the functional information present in datasets that quantify the full transcriptome.

The result thus provided the motivation to develop a laboratory method suitable for measuring the 1,000 landmark transcripts at low cost. We adopted a data-driven approach to select 1,000 landmark genes using the DS_GEO_ dataset. Because the dataset contains a non-uniform representation of various aspects of biology (for example certain tumor types such as breast and lung cancer were disproportionately represented), we applied Principal Component Analysis (PCA) as a dimensionality reduction procedure to minimize bias toward any particular lineage or cellular state. In this reduced eigenspace of 386 components (which explained 90% of the variance), cluster analysis was performed to identify tight clusters of commonly co-regulated transcripts. We applied an iterative peel-off procedure to select the centroids (Tseng and Wong, 2005). At each iteration we identified the most concordant clusters. For each cluster the transcript closest to the centroid was selected as a candidate landmark gene. All cluster members were subsequently dropped and the procedure was repeated to identify additional clusters in the remaining feature space. Transcripts nominated as landmarks through this process were then tested empirically to assess ability to measure levels accurately in the assay protocol as described below.

### L1000 assay platform

Having established through simulations that measuring ∼1,000 landmarks was sufficient to capture the majority of information encoded in genome-wide expression profiles, we next sought to develop a laboratory method capable of measuring 1,000 transcripts at low cost. For this purpose, we adapted a method involving ligation-mediated amplification (LMA) followed by capture of the amplification products on fluorescently-addressed microspheres beads (Peck et al., 2006). We extended this method to a 1,000-plex reaction (Figure 1A; protocols at clue.io/sop-L1000.pdf). Briefly, cells growing in 384-well plates were lysed and mRNA transcripts captured on oligo-dT-coated plates. cDNAs were synthesized from captured transcripts and subjected to LMA using locus-specific oligonucleotides harboring a unique 24-mer barcode sequence and a 5’ biotin label. The biotinylated LMA products were detected by hybridization to polystyrene microspheres (beads) of distinct fluorescent color, each coupled to an oligonucleotide complementary to a barcode, and then stained with streptavidin-phycoerythrin. Thus, each bead was analyzed both for its color (denoting landmark identity) and fluorescence intensity of the phycoerythrin signal (denoting landmark abundance). Because only 500 bead colors are commercially available, we devised a strategy that allows two transcripts to be identified by a single bead color (Methods and Figure 1B). The final assay, which we call L1000, contains 1,058 probes for 978 landmark transcripts and 80 control transcripts chosen for their invariant expression across cell states (see Methods). The reagent cost of the L1000 assay is approximately $2 per profile.

**Figure 1.**
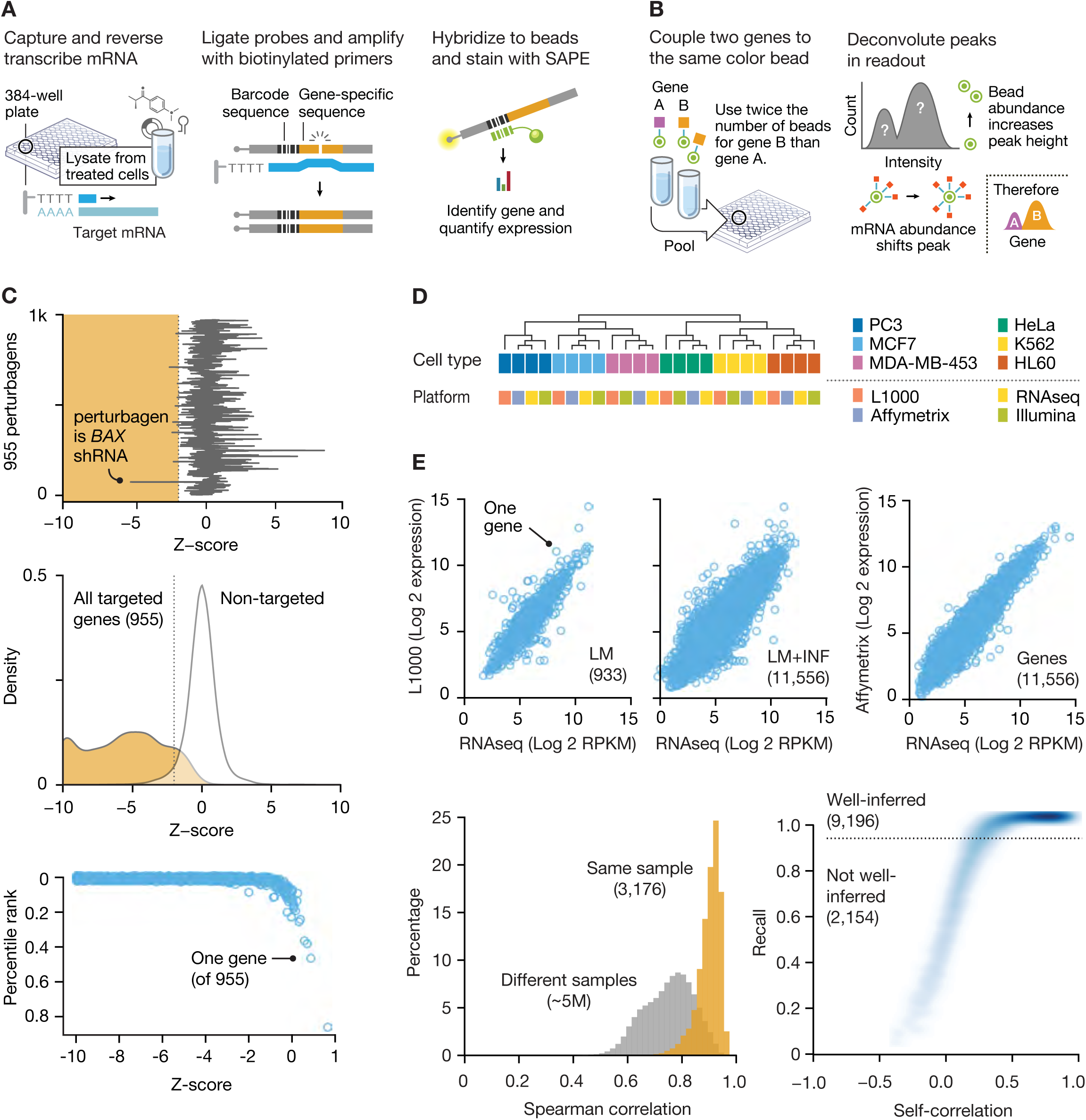
L1000 gene expression platform implementation and validation. **A. Schematic of ligation-mediated amplification (LMA) of landmark genes.** Cells growing in 384-well plates are lysed and the mRNA transcripts captured on oligo-dT-coated plates. 1,000 locus-specific oligonucleotides harboring a unique 24-mer barcode sequence are then used to perform an LMA reaction, and the biotinylated LMA products are detected by hybridization to optically addressed polystyrene microspheres (beads). Each bead is coupled to an oligonucleotide complementary to a barcode. The amount of biotinylated transcript is quantified by fluorescence after staining with streptavidin-phycoerythrin (SAPE). **B. Schematic of quantification of the ∼1,000 landmarks using 500 bead colors.** Each bead is analyzed for its bead color (denoting the landmark identity) and its phycoerythrin fluorescence intensity (denoting the landmark transcript abundance). Barcodes representing each gene are coupled to beads of the same color (one gene per bead). Genes coupled to the same color in two are combined in a ratio of 2:1 prior to use. We construct a histogram of mean fluorescent intensity, yielding a distribution that consists of two peaks, with the larger peak (by bead count) designating the expression of the gene for which double the amount of beads are present, and the smaller peak representing the other gene. Using the k-means clustering algorithm, the distribution is partitioned into two distinct components and the median expression value for each component is then assigned as the expression value of the appropriate gene. **C. Validation of L1000 probes using shRNA knockdown.** MCF7 and PC3 cells were transduced with shRNAs targeting 955 landmark genes, and generated signatures. For each targeted gene, we computed the percentile rank of its expression z-score in the experiment in which it was targeted relative to all other experiments in which it was not targeted. 841 of 955 genes (88%) rank in the top 1% and 907 of 955 (95%) rank in the top 5%, indicating the majority of L1000 probes are measured with high fidelity. Top panel: expression z-score of BAX gene in every experiment. Note it achieves the lowest expression z-score in the experiment in which it is targeted. Middle panel: Distributions of all targeted (orange) and non-targeted (white) z-scores. Bottom panel: Scatter of percentile rank versus expression z-score for 955 targeted genes. **D. Comparison of L1000 with other platforms.** Samples of purified total RNA from six human cancer cell lines were profiled on L1000, Affymetrix GeneChip HG-U133 Plus 2.0 Array, Illumina Human HT-12 v4 Expression BeadChip Array, and mRNA-seq (Illumina Hi-Seq). Data cluster by cell line and not by platform, suggesting that the cross-platform differences are smaller than the biological differences between cell lines. **E. Extended comparison of L1000 with RNA-seq and Affymetrix using patient-derived samples.** Identical RNA samples from 3,176 tissue specimens collected as part of the GTEx project were profiled on L1000 and RNA-seq. A subset was also profiled on Affymetrix microarrays. Top panels: Scatter plots of L1000 expression versus RNA-seq in landmark (left) and landmark plus inferred (middle) spaces for a single sample. L1000 correlates with RNA-seq to the same degree as Affymetrix correlates with RNA-seq (right). Bottom left: Distributions of L1000-RNA-seq correlations for the same sample (orange) and different samples (gray). This analysis yielded 3,103/3,176 (98%) with a recall (R) > 0.99 (indicating 99^th^ percentile) and all but 5 samples (99.84%) had a R > 0.95 (not shown). Bottom right:R for the L1000 inferred genes relative to their RNA-seq measured equivalents. 9,196 of 11,350 (81%) have R in the 95th percentile and are thus termed well inferred. Taken together, these results indicate that the expression profiles generated by L1000 are highly similar to their RNA-seq 0.95 equivalents.

### Optimization and validation of L1000

The fidelity of L1000 depends on its ability to quantify endogenous levels of intended landmark transcript accurately and specifically. In designing landmark-specific oligonucleotide probes, we followed several computational procedures that maximized matches to the target DNA sequence while minimizing non-specific hybridization (Wetmur, 1991) (See Methods). As sequence-based probe-selection methods are imperfect, we further optimized the accuracy of L1000 probes experimentally. Previously validated shRNAs targeting landmark transcripts were used to infect MCF7 and PC3 cells, and the L1000 platform was used to measure changes in landmark transcript abundance (Figure 1C). L1000 probes that failed to detect the perturbed transcript were re-designed. Several cycles of iteration resulted in a final L1000 probe set comprising 978 landmarks and the 80 control genes (Table S2).

Having chosen the L1000 landmark transcripts using an entirely data-driven approach optimized for maximal information rather than on biological function, we asked whether the landmarks are enriched in any particular functional classes (e.g., transcription factors). We computed hypergeometric overlap statistics and found no substantial enrichment for any particular protein class. Similarly, we found no evidence of developmental lineage bias, based on an analysis of landmark expression patterns across 30 tissue types (Figure S1B). These results are consistent with our goal of identifying landmark transcripts based on their ability to faithfully reconstruct the transcriptome, without over-optimizing for any particular pathway, cell type or cell state.

### L1000 reproducibility

As an initial measure of L1000 reproducibility, we analyzed technical replicates of 6 cancer cell lines in which aliquots of the same RNA sample were subjected to replicate L1000 profiling (12 replicates in each of 3 batches, yielding 36 replicates per cell line). Within each cell line, we computed the Spearman correlation between all pairwise combinations of replicates, excluding the comparison of each replicate to itself. We found that nearly all pairwise comparisons achieved a high correlation (>0.9) suggesting low sample-to-sample variability (Figure S1C). Furthermore, intra-batch variation was comparable to inter-batch variation. Overall, analysis of 216 such replicates resulted in a median correlation coefficient of 0.95, indicating extremely high technical reproducibility of the assay.

### Comparison of L1000 to RNA-seq

RNA sequencing (RNA-seq) has become the standard for gene expression profiling, and thus we sought to benchmark L1000 against it. We note that while RNA-seq is attractive given its unbiased nature, it suffers from technical complexity in library preparation, and most importantly, inability to detect non-abundant transcripts without deep sequencing that results in higher costs. The L1000 platform, like DNA microarrays, is hybridization-based, thus making the detection of non-abundant transcripts feasible. As an initial assessment of cross-platform performance, mRNA samples from 6 cell lines were profiled on L1000, Affymetrix U133A and Illumina BeadChip arrays, and by RNA-seq. Hierarchical clustering of these data grouped samples by cell type, not measurement platform (Figure 1D), and moreover cross-platform gene expression correlation was at the transcript level (Figure 1E, upper panel).

To more extensively compare L1000 to RNA-seq, we analyzed 3,176 samples that were profiled on both platforms. The RNA samples, representing 30 tissues were previously sequenced as part of the GTEx Consortium (The GTEx Consortium, 2015); DS_GTEx-rnaseq_), and we subjected aliquots of those same samples to L1000 profiling (DS_GTEx-L1000_). We then computed sample self-correlations for the 3,176 samples and the median sample self-correlation was 0.84, with a notably right-shifted distribution relative to non-self correlations (Figure 1E, lower panel left). We also measured sample *Recall* (R_sample_, see Methods), wherein a given L1000 profile is forced to compete with all other RNA-seq profiles in order to find its RNA-seq counterpart. This analysis yielded 3,103/3,176 samples (98%) with a R_sample_ > 0.99 (indicating 99^th^ percentile) and all but 5 (99.84%) had a R_sample_ > 0.95 (Figure S1D). Taken together, these results indicate a strong degree of similarity in profiles across L1000 and RNA-seq platforms.

### Inferring gene expression from L1000 landmarks

Using 8,555 RNA-seq samples (Dataset DS_GTEx-rnaseq_) as an independent test set, we used landmark transcript measurements to infer the remainder of the transcriptome. As a test of inference accuracy, we analyzed gene-level recall (R_gene_) for each of the inferred genes and assessed performance by comparing the result to a null distribution of correlations between all inferred transcripts and all measured transcripts. This analysis showed that inference was accurate (defined as R_gene_ > 0.95) for 9,196 of the 11,350 inferred genes (81%). When combined with the 978 measured landmarks, the L1000 platform thus measures or infers with high fidelity 83% of transcripts, but yields poor inference for 17% (Figure 1E, lower panel right and Table S3). Inferences for these 17% were therefore not used in any of the analyses that follow.

### Generation of the first million L1000 Connectivity Map profiles

Having validated a low-cost, high-throughput L1000 assay, we set out to create a large-scale Connectivity Map dataset as a community resource. We expanded on the Connectivity Map pilot dataset in several dimensions.

First, we increased the number of small molecule perturbations from 164 FDA-approved drugs to 19,811 small molecule drugs, tool compounds and screening library compounds including those with clinical utility, known mechanism of action, or nomination from the NIH Molecular Libraries Program. Each compound was profiled in triplicate, either at 6 or 24 hours following treatment.

Second, we expanded in the dimension of genetic perturbation by knocking down and overexpressing 4,372 genes selected on the basis of their association with human disease or their membership in biological processes or pathways. Each genetic perturbation was profiled in triplicate, 96 hours after infection. For overexpression studies, a single cDNA clone representing the entire coding sequence was lentivirally transduced (Moffat et al., 2006). For loss-of-function experiments, three distinct shRNAs targeting each gene were profiled.

Third, we expanded in the dimension of cell lines. Whereas the CMap pilot dataset contained profiles in just three cancer cell lines, we expanded the dataset to include as many as 77 cell lines. Well-annotated genetic and small molecule perturbagens were profiled in a core set of 9 cell lines, yielding a reference dataset we refer to as *Touchstone v1*. Uncharacterized small molecules without known mechanism of action (MOA) were profiled variably across 3 to 77 cell lines, yielding a dataset we refer as *Discovery v1*. Details of the contents of both datasets are available in Table S4.

In total, we generated 1,328,098 L1000 profiles from 42,553 perturbagens (19,811 small molecule compounds, 18,493 shRNAs, 3,627 cDNAs, and 622 biologics) for a total of 476,251 signatures (consolidating replicates), representing over a 1,000-fold increase over the CMap pilot dataset. We term this first release of an L1000-based compendium as *CMap-L1000v1* (Figure 2A). All data, at multiple levels of pre-processing are available via GEO (accession GSE92742 and pre-processing code via GitHub), and for easier use via the CLUE analysis environment (https://clue.io; see below and Figure 2B).

**Figure 2.**
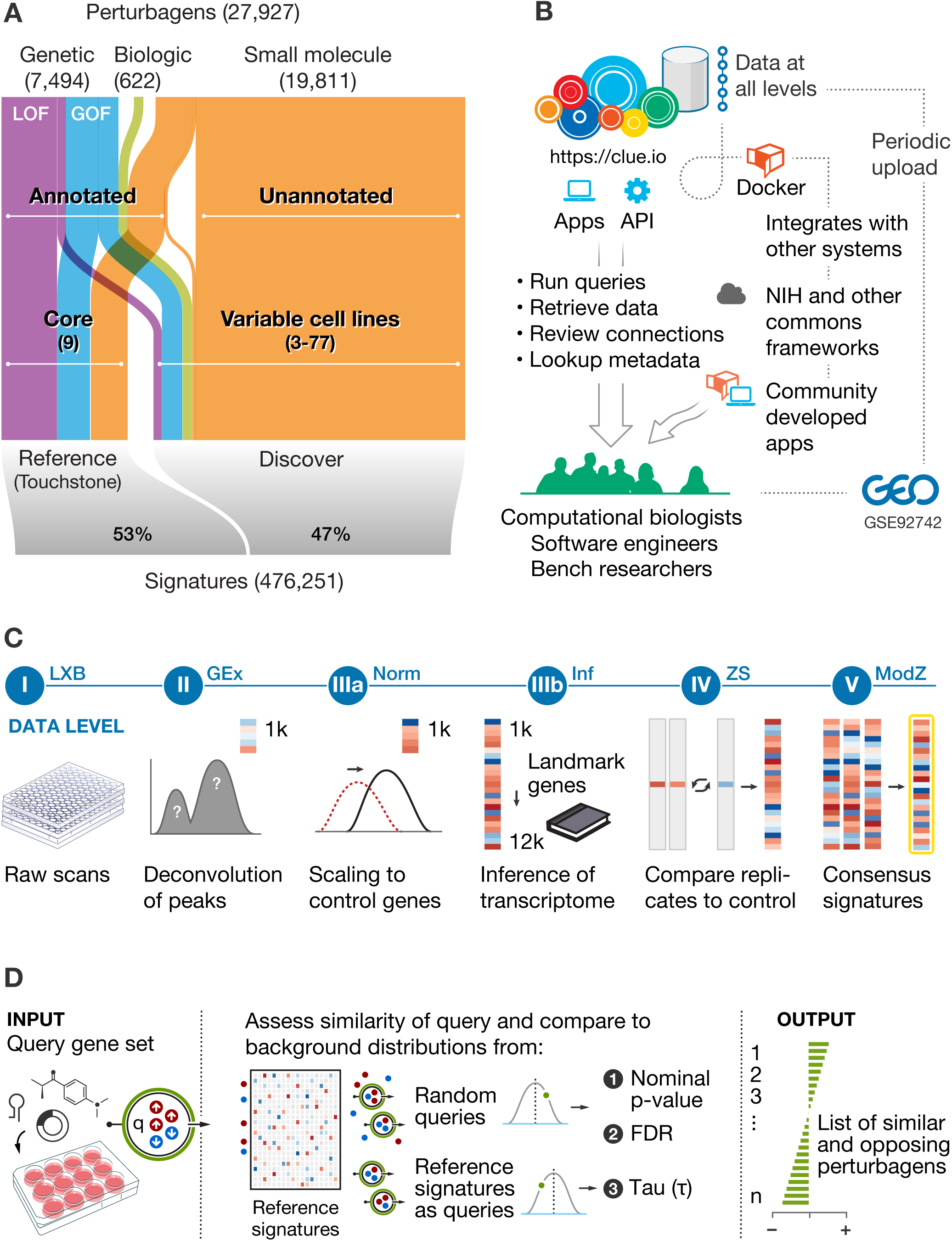
L1000 dataset coverage, signature generation, and data access. **A. Depiction of data in *CMap-L1000v1***. Perturbations are divided by type, annotation status, and cell line coverage. The number of genetic perturbagens refers to the number of unique genes targeted by either shRNA or over-expression. Annotated perturbations profiled systematically across 9 core cell lines comprise the Touchstone portion of the dataset (CMap-TS) and are useful for reference comparisons, while unannotated reagents comprise the Discover portion and present opportunities for discovery. **B. Modes of access to analysis tools and data via the clue.io software platform.** The clue.io platform enables computational biologists, bench scientists, and software engineers to leverage CMap by offering web applications for analysis, APIs and docker containers for code and data access. Raw data can be downloaded from NCBI GEO. **C. Schematic of signature generation and data levels.** 1) Raw bead count and fluorescence intensity are acquired from Luminex scanners 2) Data are deconvoluted to assign expression levels to two genes measured on the same analyte 3a) Data are normalized to adjust for non-biological variation 3b) 12,328 genes are inferred from the 978 landmarks 4) Data are converted to differential expression values 5) Replicate profiles are collapsed into signatures. **D. Schematic of query analysis.** A query is specified by lists of up-and down-regulated genes. The similarity between query and all signatures in CMap-TS are computed. Normalized similarities are converted to p-value and FDR, and tau via comparison with compendia random and reference queries, respectively. Finally, perturbagens are sorted by tau to provide the list of most similar and opposing perturbagens, thus providing hypotheses for potential follow-up.

### CMap query methodology

The connectivity workflow involves interrogating the CMap database of *signatures* with a *query* (a set of differentially expressed genes representing a biological state of interest). Each of the signatures in the database represents a weighted average across the 3 biological replicate perturbations (see Methods). This moderated z-score procedure serves to mitigate the effects of uncorrelated or outlier replicates, and can be thought of as a ′de-noised′ representation of perturbational response (Figure 2C). The similarity of the query to each of the CMap signatures is computed, thus yielding a rank ordered list of the 473,647 signatures in the CMap-L1000v1 dataset. However, simply sorting by degree of similarity can be misleading because such a procedure alone does not statistically address issues such as magnitude of gene expression change or specificity of observed connections.

We therefore developed a Connectivity Score (Figure 2D) that provides three measures of confidence: 1) a nominal p-value derived by comparing the similarity between the query and reference signature, using the Kolmogorov-Smirnov enrichment statistic (Subramanian et al., 2005), to a null distribution of random queries;

2) a false discovery rate (FDR) that adjusts the p-value to account for multiple hypothesis testing given the large numbers of comparisons in the dataset; and 3) Tau (τ), which compares an observed enrichment score to all others in the database. τ is a standardized measure ranging from -100 to 100 and can be used to compare results across queries; a τ of 90 indicates that only 10% of perturbations showed stronger connectivity to the query. A connection with a significant p-value and FDR but low τ would suggest a highly promiscuous perturbagen whose connections are not unique.

These Connectivity Score metrics thus constitute a statistical framework that provides a holistic quantification of the relationship between a query and a perturbagen, as opposed to merely sorting by degree of similarity, as was used in the CMap pilot. Additionally, while the Connectivity Scores are generated on each cell type individually, we summarize those scores across all profiled cell types and thus provide a measure of robustness. Importantly, this analytical approach is platform-independent, allowing users to create query signatures from any gene expression platform.

### Feasibility of querying a million-profile compendium

With an analytical framework in hand, we next set out to test the CMap compendium for its ability to produce biologically meaningful connections. That is, while our analysis of replicate measurements demonstrated that L1000 is robust, it is conceivable that as the size of the dataset increased, so might biological and technical noise, thereby obscuring real signal. Additionally, laboratory and platform batch effects might overwhelm true signal thereby making it harder to find genetic or pharmacological connections when query signatures are generated by other labs on other expression platforms. To address this, we compiled a database of 7,578 perturbational signatures from public sources from which we identified 1,143 perturbational profiles that matched a *CMap-L1000v1* perturbagen, and were therefore eligible for Recall analysis. For each query, we assessed whether it connected to its equivalent in *CMap-L1000v1* at a high level of confidence (defined as NP <=0.05, FDR <= 0.25 and |τ| >= 90). 909/1,143 queries (80%) exhibited the expected connectivity. We note that the inference of expression values from landmarks was essential for the success in recovering connections. 20% of connections were lost when the analysis was restricted to landmarks only. Furthermore, 48 query signatures contained zero landmark transcripts and were therefore not analyzable without inference of the remainder of the transcriptome. Overall, we conclude that the Connectivity Map dataset has a high degree of biological coherence that can be used to discover connections to user-defined, external queries (see Methods and Table S5 for queries tested and scores).

### Discovering off-target effects of shRNAs

The scope of the L1000 dataset provides an unprecedented opportunity to examine the biological effects of RNA interference, in particular, the off-target effects of shRNAs. Such off-target effects have long been suspected, but there has been no way to systematically quantify the magnitude and nature of a given shRNA’s on-target versus off-target gene expression effect. To address this, we analyzed 13,187 shRNAs targeting 3,799 genes (3 shRNAs/gene) across 9 cell lines. We then compared each pair of shRNA-induced L1000 profiles focusing on the degree of similarity between two shRNAs targeting the same gene (“shared gene”) and two shRNAs targeting different genes but sharing the same 2-8 nucleotide seed sequence known to contribute to off-target effects (“shared seed”) (Jackson et al., 2003). Figure 3A shows that as a group, shared gene similarity is only slightly greater than that observed between randomly selected pairs of shRNAs. In contrast, shared seed pairs were dramatically more similar compared to the null distribution, indicating that the magnitude of off-target effects of shRNAs substantially exceeds the magnitude of their on-target effect. To address the strong off-target effects of shRNAs, we reasoned that while on-target gene expression effects of different shRNAs targeting the same gene should be the same, their off-target effects should not. We therefore developed an algorithm to produce a Consensus Gene Signature (CGS) that reflects the consistent (and therefore on-target) gene expression effects of shRNAs and used the CGS output for all analyses that follow. The CGS method and its validation are being reported elsewhere. Examples of the CGS recovering compound-gene connections that were missed at the individual shRNA level are shown in Figure 3B.

**Figure 3.**
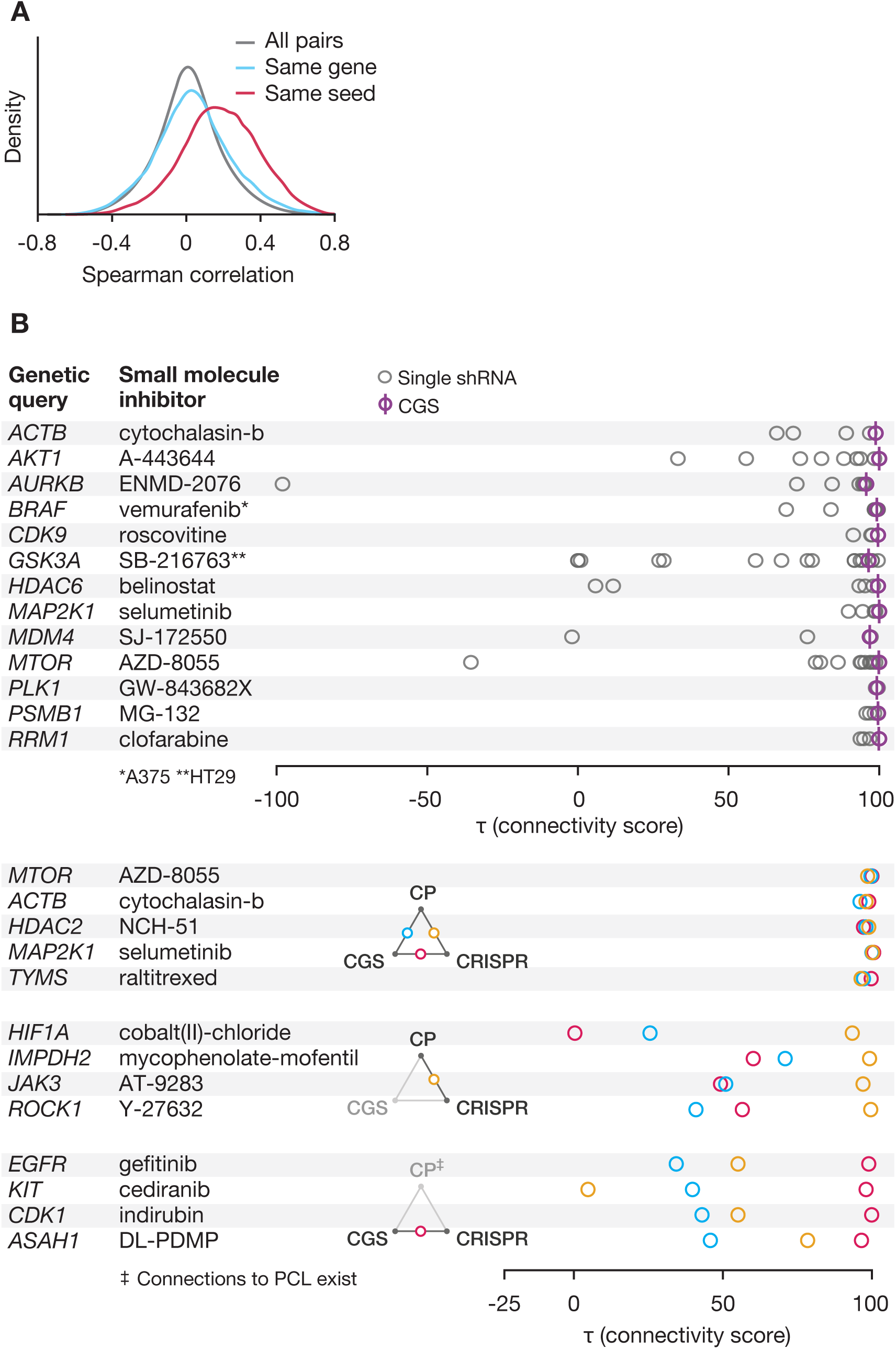
Analysis of genetic loss of function perturbations. **A. Notable off-target effects of shRNAs.** Distributions of spearman correlations between signatures of 12,961 shRNAs targeting the same gene but having different seed sequences (blue), targeting different genes but having the same seed (red) and all pairs of shRNAs (gray). 12,691 shRNAs targeting 3,724 unique genes were considered and data shown is for the A549 cell line. The all pairs distribution was randomly down-sampled to 10M points for plotting. While shRNAs targeting the same gene are better correlated than all pairs, shRNAs with the same seed are much more correlated, suggesting that the shRNA seed effect may dominate the gene effect, presenting a problem for analysing shRNA signatures. Data is shown from the A549 cell line. **B. Consensus Gene Signature (CGS) improves on-target signal.** A consensus gene signature (CGS) is computed from a weighted average of signatures of independent shRNAs targeting the gene. Connectivity to annotated small molecules targeting each gene is markedly improved by CGS over individual shRNAs, suggesting that the CGS procedure mitigates the seed effect inherent to individual shRNAs and enhances on-target signal. **C. CRISPR knockout augments compound-target analysis.** Top: Consistency between Loss of Function (LoF) signatures from CRISPR and CGS enhances confidence in connectivity to small molecules. Middle: CRISPR-based LoF recovers some connections to small molecules missed by CGS. Bottom: Lack of compound-target connectivity, despite consistency between LoF reagents and validated compound signature suggests non-equivalency of genetic and pharmacological agent derived signatures.

### Characterizing small molecule function

A theoretical feature of a large-scale Connectivity Map is the ability to determine mechanism of action (MOA) of a small molecule, based simply on similarity of gene expression induced by genetic perturbagens or compounds of known function. As a first step in assessing feasibility of such an approach, we sought to determine whether known MOAs of drugs and tool compounds could be recovered by the CMap. This is challenging, however, because most compounds lack ground truth with respect to their protein targets and associated pathways. While certain targets of many compounds are known, even those compounds may have additional unidentified targets, and for some compounds, precise MOA remains unknown. Nevertheless, we used multiple information resources to associate 1,902 compounds to protein targets and associated pathway members profiled in the CMap. This led to 58,820 expected relationships that could plausibly be recovered in the *CMap-L1000v1* compendium (see Methods and screening library publication for details of annotations) (Corsello et al., 2017). We then sought to recover those relationships from among the approximately 160 million pairwise relationships (connections) that could be assessed across the compendium.

For each compound, we computed the true positive rate (i.e., recovery of expected relationships). We refer to these expected relationships as *expected pairs*. To estimate the false positive rate, we counted the connections between compounds and genetic or pharmacologic perturbagens annotated as having a relationship with a different small molecule in the dataset. We refer to such relationships as *null pairs*. We then plotted the true positive rate against the false positive rate at various thresholds of statistical significance, thereby generating an ROC curve from which an AUC could be calculated. An AUC >0.6 is typically regarded as signifying a positive signal. At that cut-off, an average of 45% of expected relationships were recovered in any one of the 9 cell lines tested (range 29%-58% per cell line). This number rose to 63% when Connectivity Scores were summarized across all 9 lines (full results in Table S6).

While this result is encouraging, 37% of compounds did not show evidence of connection to their expected targets. Failure to recover such connections could be explained by many factors including i) incomplete inhibition of the target by the compound, ii) off-target effects of compounds and genetic perturbations, iii) missing information in the L1000 read-out, iv) incorrect literature-based annotations of compounds, v) biological differences between small molecule inhibition of specific aspects of protein function (e.g., enzymatic inhibition) compared to complete loss of function (e.g., scaffolding functions) induced by shRNA-mediated knock-down, and vi) the existence of previously unrecognized *bona fide* connections that effectively penalize the known connections – particularly if the novel connections are stronger than the expected ones.

An even more stringent approach is to ask whether the direct protein target (not just the affected pathway) of a small molecule could be recovered by the CMap. For this analysis, we evaluated 926 drugs targeting 283 proteins, and found that 51% of these connections were recovered (defined by being in the top 100 gene perturbations, representing the top 3% of genes tested, Table S7). We also explored whether failure to recover expected connections might be explained at least in part by cellular context. To address this, we selected 25 compounds lacking strong connections to their expected direct targets in the 9 core cell types, and we re-profiled them in additional 39 cell lines. In 12/25 cases (48%), the expected connections were revealed (Table S7, first 12 rows). The rescue of expected connections by expanding the number of cell lines profiled could be explained by either differences in cell state (e.g., developmental lineage) or by genotype (e.g., mutation status). We note that CMap connections can indeed be sensitive to genotype. For example, we observed that the signature of the MDM2 inhibitor AMG-232 differed in MCF10A breast epithelial cells that were either *TP53* wild-type or homozygously deleted, reflecting MDM2’s role as a negative regulator of TP53 (Figure S2A).

Taken together, these results indicate that the CMap can serve as a powerful strategy to assign mechanism of action to small molecules, and furthermore suggests that the success of this approach will improve by further expansion of the CMap to include maximally genetically and developmentally diverse cell types.

### Defining perturbagen classes (PCLs)

The analytical strategy described in the sections above establishes a statistically principled method for ranking the significance of connections to a CMap query. A challenge, however, is that the analysis returns a rank-ordered list of connections, leaving the user to extract biological meaning from the list (e.g., by observing similar compounds or pathway-related genes at the top of the list). In some cases, such biological insights are obvious, but often they are not. We reasoned that observed connections could be analyzed in a manner analogous to defining consensus signatures across multiple shRNAs targeting the same gene. That is, while any given member of an MOA class would likely have a multitude of targets, integrating signatures across several examples of an MOA class would sharpen the on-target signal, while diminishing off-target effects, as has been proposed previously (Seashore-Ludlow et al., 2015).

We codified this class-level annotation by first identifying groups of compounds of distinct chemical structure that share the same MOA. We also identified groups of genetic perturbagens belonging to the same gene family or being commonly targeted by the same compounds. These perturbagen classes (PCLs) were then further refined by excluding from the PCL any compounds that failed to empirically connect with their cognate class members based on L1000 connectivity analysis (see Methods and Figure 4A). This procedure yielded 171 high confidence genetic and pharmacologic PCLs (Table S8). Observing strong connectivity to a PCL, as opposed to an individual perturbation, thus provides users with a higher level of confidence in interpreting biological mechanism from CMap analyses.

**Figure 4.**
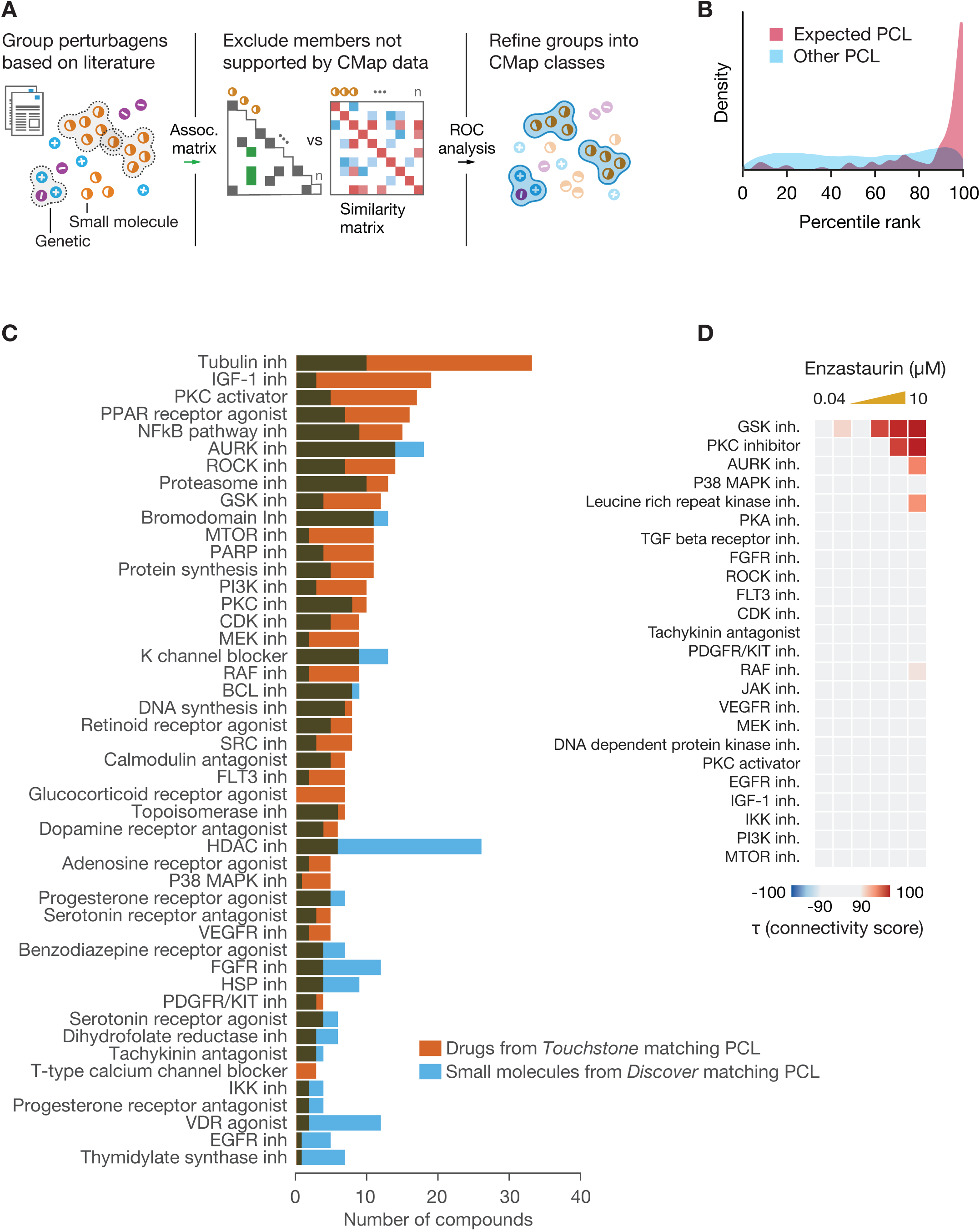
Reference perturbagen classes for CMap discovery. **A. Schematic of Perturbagen Class (PCL) definition.** Reference PCLs are established by integrating existing drug and gene annotations with CMap connectivity results. Left: Annotations are first gathered from literature sources and perturbagens with shared annotations are grouped to form candidate sets. Middle: Perturbagens that do not sufficiently recover their expected connections via ROC analysis, are excluded. Right. The remaining members are then assessed for intra-group connectivity and groups that are sufficiently interconnected are retained as PCLs. These PCLs serve as robust sensors of biological activities and are useful in interpreting CMap query results. **B. PCL validation.** 137 compounds with known activities corresponding to one or more of 54 PCLs, but that were not used in PCL construction, were prospectively profiled across multiple cell types and subjected to PCL connectivity analysis. The histogram shows the rank of each expected PCL connection for these 137 compounds (red) versus the rank of all unexpected PCL connections (blue). The expected PCL distribution is notably right-shifted, indicating that PCLs are accurately reading out the expected activities from an independent compound set. **C. Using PCLs for discovery.** 3,333 known drugs and 2,418 unannotated but transcriptionally active compounds were subject to PCL connectivity analysis. The plot shows the count of strong and selective connections to validated PCLs by known drugs not annotated with the given PCL’s activity (blue) and unannotated compounds (orange). The novel connections of known drugs present hypotheses for secondary mechanisms and/or off-target effects, while, for unannotated compounds, they may suggest the molecule’s primary activity. **D. Detecting multiple drug activities using PCLs.** The known PKC inhibitor enzastaurin was profiled in CMap across multiple doses. Connectivity to each established kinase inhibitor PCL is shown in the heatmap. Strong dose-responsive connections were observed to the PKC and GSK3 inhibitor PCLs.

To test this hypothesis, we profiled 137 test compounds known to share a mechanism with one or more of 54 small molecule PCLs, but which were not used in the construction of the PCL. L1000 profiling indeed revealed that for 41/54 classes (76%), the test compounds connected to their designated PCL in multiple cell types (Figure 4B). For an additional 7/54 (13%), a selective connection was observed in a single cell type. The remaining 6/54 (11%) did not reconnect at a threshold of τ >90.

We next asked whether drugs with established MOA also had unexpected, strong, and selective connections to a validated PCL. Selectivity was defined as the fraction of PCLs to which a drug failed to connect at a given τ threshold. 132 drugs (3.9%) had such off-target connections (See Methods and interactive matrix at https://clue.io). For example, compounds showing connectivity to the protein kinase C (PKC) inhibitor PCL were often also strongly correlated to the GSK3 inhibitor PCL. 44 such dually connected compounds were found (τ>=95, selectivity >=0.85), including the PKC inhibitor enzastaurin which showed dose-responsive connectivity to both PKC and GSK3 inhibitor classes (τ_GSK_=99.79 τ_PKC_=99.47, selectivity=0.88) across a similar dose range (Figure 4D). Interestingly, synergy between compounds targeting these pathways has been reported (Rovedo et al., 2011), and the Kinomescan biochemical binding assay confirms that enzastaurin is indeed also a potent GSK3 inhibitor with a biochemical K_D_ of 8 nM (Davis et al., 2011). Importantly, however, not all GSK inhibitory compounds have PKC off-target effects. 23 small molecules (Table S9) showed GSK PCL selectivity. Interestingly, the top scoring GSK3 PCL connection without strong PKC connectivity (τ_GSK_=96.66, τ_PKC_=13.71, selectivity=0.95) was the compound JNJ-16259685, developed against the metabotropic glutamate receptor GRM1 (also known as mGluR1). Interestingly, GRM1 has been recently reported to be an upstream effector of GSK3β (Liu et al., 2005). These results highlights the use of the Connectivity Map to identify compounds with a desired level of selectivity, and shows that even well-characterized compounds may have unexpectedly strong direct or downstream effects.

Importantly, compounds connected to multiple PCLs were not limited to kinase inhibitors. For example, examination of the BET bromodomain inhibitor PCL recovered compounds developed as BET inhibitors (JQ1-(+), I-BET-151, I-BET-762, PFI-1, SB-203580 and LY-303511 not used in PCL definition) but also compounds developed against other targets (Table S10). Examples include alprazolam, a benzodiazepine shown to also interact with BET proteins (Filippakopoulos et al., 2012), XMD11-85H, a BRSK2 kinase inhibitor, droxinostat, an HDAC inhibitor, and BRD-1103, a tool compound inhibiting JAK3. Followup of XMD11-85H, droxinostat and BRD-1103 via bromoscan revealed inhibitory activity against one or more of the 32 bromodomains tested (Figure S3A).

We note that in the future, as the number of compounds in the Connectivity Map grows, splitting of current PCLs to reflect subclasses with distinct patterns of selectivity may be possible. For example, the histone deacetylase (HDAC) inhibitor PCL class currently has 20 members, each with varying selectivity against the 13 HDAC proteins. Clustering the L1000 gene expression data revealed clear substructure within the PCL, with pan-HDAC-inhibitory compounds forming a distinct cluster, and compounds selective for either HDAC6 or HDAC1,3 and 8 forming distinct clusters (Figure 5A).

**Figure 5.**
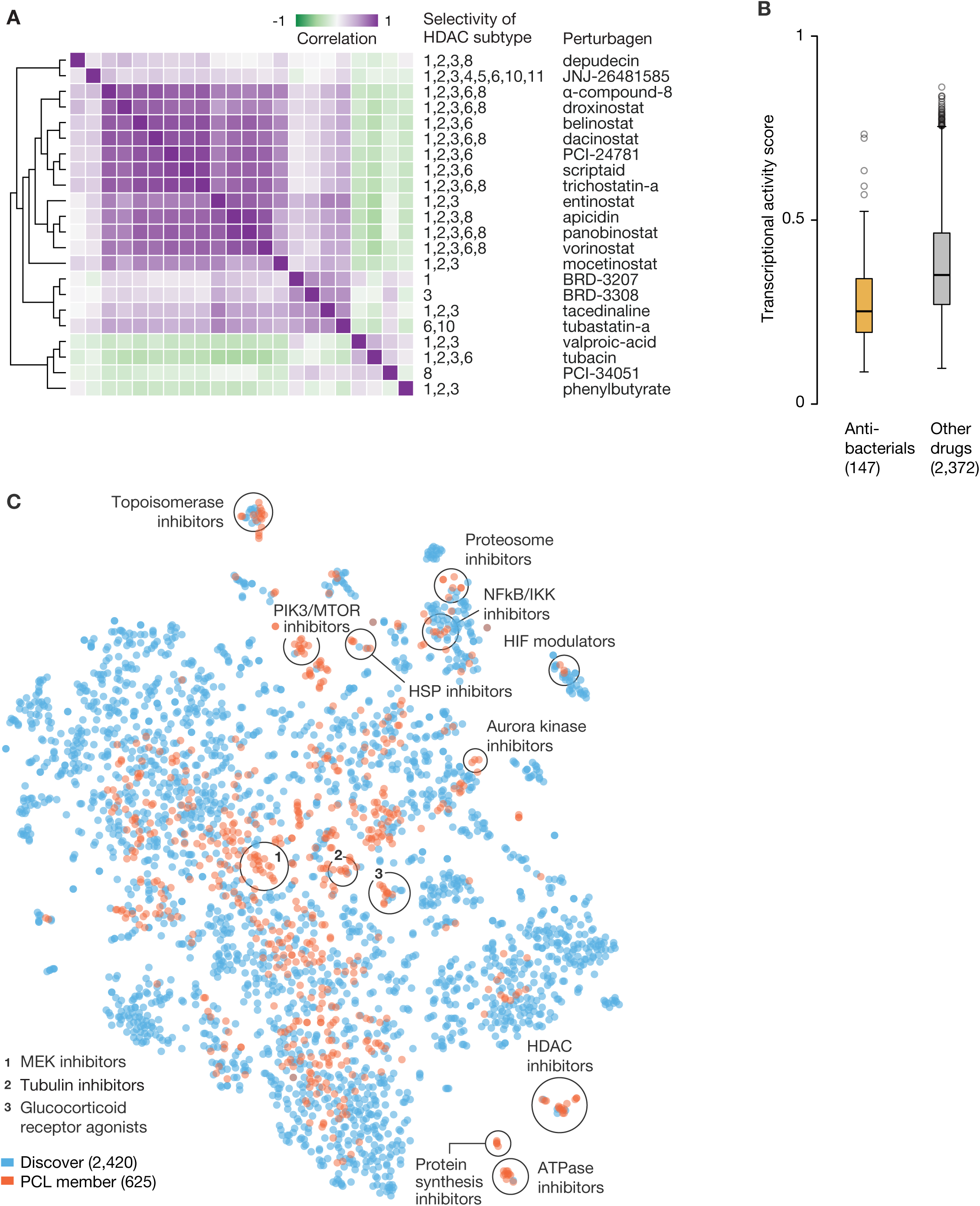
Characterizing known and unexpected activities of small molecules. **A. Antibacterials exhibit lower transcriptional activity than other drugs.** Distributions of the maximum TAS per compound for 147 antibacterials and 2,372 known drugs in CMap-TS. The antibacterials’ TAS distribution is significantly lower than that of other drugs, which matches the intuition that these compounds should generally have lower magnitude effects in human cells. (p-value corresponds to 2-sided KS test using R’s ks.test function). **B. Comparison of unannotated compounds with known drugs.** The figure shows a tSNE projection of the signatures of 2,418 unannotated but transcriptionally active compounds (orange) along with PCL members (blue). There are several highlighted instances of unannotated compounds clustering with drugs of the same mechanism, suggesting hypotheses for the unannotated compounds. In addition, the unannotated compounds occupy regions not covered by known drugs, presenting opportunities for novel mechanism discovery and potential expansion of CMap-TS **C. HDAC inhibitor PCL substructure.** Hierarchical clustering of pairwise connectivities of the HDAC inhibitor PCL members reveals substructure within the class. The pan-HDAC inhibitors generally cluster together, distinct from the more isoform-selective compounds, suggesting that gene expression can be used to further stratify compounds within the same class.

Taken together, these results suggest that Connectivity Map analysis can be used to ascertain the selectivity of drug candidates by profiling the breadth of their cellular effects in living cells. In most cases, lack of selectivity is considered a liability. In principle, multi-targeted compounds could have desirable therapeutic benefits, and the CMap could be used to guide medicinal chemistry toward such dual activities. We note that such an approach is particularly powerful when considering protein targets for which family-wide profiling methods do not exist.

### Cellular context

To study the effect of cellular context on perturbational responses, we compared the signatures of 2,429 drugs across a panel of 9 cancer cell lines. On average 38% of compounds scored as transcriptionally active (based on signature strength and replicate consistency) in any single cell type (range 28%-45%) and 92% of small molecule drugs scored as active in at least one cell line. Of 1,399 (58%) compounds active in at least 3 cell lines, 26% (corresponding to 15% of all compounds) produced highly similar signatures across the entire panel, whereas perhaps not surprisingly, the remainder were active in only 1 or 2 cell lines or produced a diversity of cellular signatures (see Methods and Figure S2B, S2C).

As might be expected, connections with support across multiple cell types tended to target core cellular processes (e.g., ribosomal function, proteasome complex), whereas compounds with reproducibly cell-type-selective patterns of connectivity tended to target more specialized mechanisms. For example, connectivity between multiple glucocorticoid receptor agonists was restricted to those cell types in which the glucocorticoid receptor gene *NR3C1* was expressed (Figure S2D, upper panel). Connectivity between multiple small molecule PPARG agonists was greatest in HT29 and PC3, the two core cell lines with the highest baseline expression of PPARG itself (Figure S2D, lower panel). Similarly, the connection between androgen receptor (*AR*) knockdown and the AR antagonist nilutamide was strongest in the AR-expressing cell line VCAP (Figure S2E). We also note that the naturally occurring genetic diversity of disparate cell lines adds to the value of CMap analyses. For example, connections between genetic perturbation of the MAP kinase pathway and small molecule inhibitors of RAF or MEK kinases were strongest in the cell lines A375 and HT29 that harbor BRAF V600E kinase-activating mutations. This and other additional examples of lineage-selective connections are shown in Figure S2E.

### Identifying bioactive subsets of small molecule screening libraries

Modern methods in chemical synthesis now make it possible to create large numbers of structurally diverse small molecule compounds. However, many such compounds fail to engage specific protein targets or to even enter living cells. We asked whether an L1000 profile could serve as a sensor for biological activity. If so, screening chemical libraries with L1000 might enable rapid elimination of compounds lacking obvious activity and help prioritize others for subsequent cell-based screening. In support of this notion, and consistent with our earlier studies (Wawer et al., 2014), we found that whereas 2,232/2,429 (92%) established drugs yielded a strong L1000 transcriptional response (defined as Transcriptional Activity Score (TAS) >0.2; see Methods), only 2,418/16,527 (15%) un-optimized compounds had high TAS scores. We note, however, that compounds with cell-type selective bioactivity might be missed by this approach.

Interestingly, the TAS-low drugs were enriched in antimicrobial agents that would not be expected to target human proteins (Figure 5B). An exception to this finding is the antimicrobial triclosan (used in consumer products), which yielded a particularly high TAS score, consistent with its having effects in mammalian cells. We note that the safety of triclosan has recently been questioned (Dinwiddie et al., 2014; Yueh et al., 2014). Projection of TAS-high compounds in two dimensions using the t-SNE method (Van der Maaten and Hinton, 2008) shows that many uncharacterized compounds cluster with existing PCLs (Figure 5C). A more quantitative approach to assigning MOA of uncharacterized compounds is described below.

### Discovery of MOA of novel small molecules

Having demonstrated the ability to recover *expected* small molecule-protein target connections, we next asked whether the Connectivity Map could be used as a means to identify the MOA of previously uncharacterized compounds. For this analysis, we queried the Touchstone reference database with signatures from L1000 profiling of compounds from various screening libraries.

We put particular attention to the discovery of novel kinase inhibitors simply because of the availability of facile methods for validating CMap predictions. For example, our analysis indicated that the unannotated compound BRD-2751 showed strong connectivity to the Rho-associated protein kinase (ROCK) PCL, suggesting that it might in fact be a ROCK inhibitor. To test this hypothesis, we subjected the compound to kinome-wide binding measurements (using the Kinomescan assay) and found that precisely as predicted, the compound has a KD of 56 nM against ROCK1 (Figure 6A). We note that while the compound had not been previously reported to be a ROCK1 inhibitor, its chemical structure is reminiscent of canonical ROCK inhibitory compounds. As another example, several compounds (BRD-5161, BRD-5657, and BRD-9186) were predicted to function as MTOR and/or PI3 kinase inhibitors. Kinomescan dose-response profiling confirmed that the three compounds were indeed MTOR/PI3K inhibitors, spanning a range of potencies and selectivities (Figure S3B). As a large number of potent and selective PI3K/MTOR inhibitors already exist, we did not investigate these novel compounds further. However, these results illustrate the power of looking up MOA information directly from the Connectivity Map.

**Figure 6.**
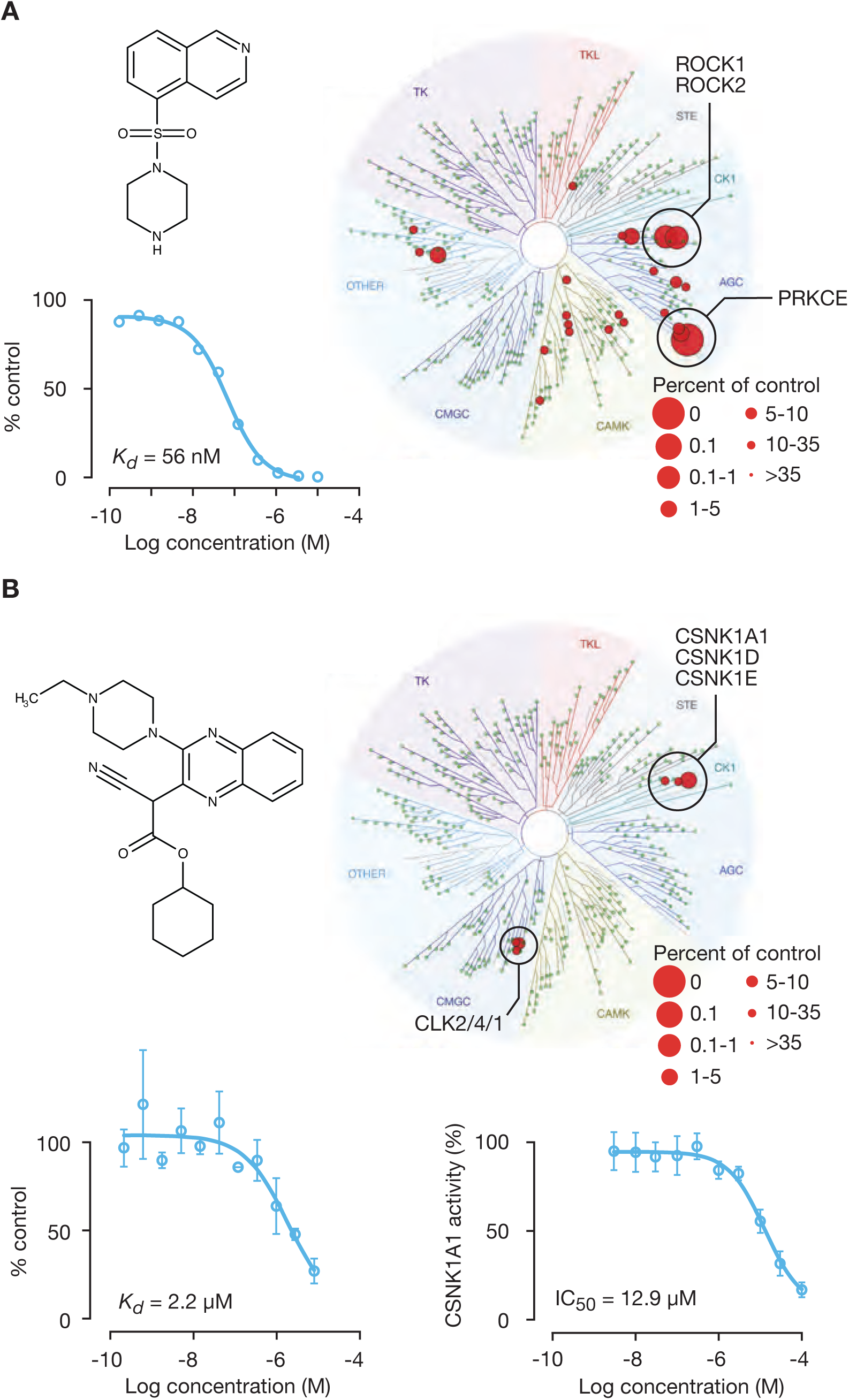
Kinase inhibitor discovery using reference transcriptional signatures. **A. Discovery of a potent ROCK1/ROCK2 inhibitor.** Top left panel: The chemical structure of BRD-2751, a predicted ROCK inhibitor is shown. Right: TREEspot selectivity profile of Kinomescan binding assay confirmed compound binding to ROCK1/ROCK2. Bottom left: Dose response testing by Kinomescan showed ROCK1 Kd of 56 nM. **B. Discovery of a highly specific CSNK1A1 inhibitor**. Top left panel: The chemical structure of BRD-1868, a predicted CSNK1A1 inhibitor, is shown. Top right: TREEspot image of Kinomescan binding assay performed with BRD-1868 at 10 micromolar demonstrated inhibition of only 6 out of 456 kinases tested including CSNK1A1. Bottom left: CSNK1A1 binding by BRD-1868 was confirmed by Kinomescan with Kd of 2.2 uM. Error bars indicate standard deviation between technical replicates. Bottom right: BRD-1868 inhibits phosphorylation of peptide substrate by CSNK1A1 with IC50 of 12.9 uM. Error bars indicate standard deviation between technical replicates. Kinase selectivity images generated using TREEspot Software Tool and reprinted with permission from KINOMEscan, a division of DiscoveRx Corporation.

### Discovery of a selective CSNK1A1 inhibitor

We next asked whether we could use the Connectivity Map to discover a compound with a particular activity – in this case, an inhibitor of an emerging therapeutic target in cancer: Casein Kinase 1A1 (CSNK1A1). CSNK1A1 is a serine-threonine kinase that was recently reported as an essential gene in certain subtypes of myelodysplastic syndrome and acute myeloid leukemia, and also has been recently shown to be targeted for degradation by the drug lenalidomide, which is particularly effective in MDS patients with chromosome 5q deletion (the locus of the *CSNK1A1* gene) (Järås et al., 2014; Krönke et al., 2015; Schneider et al., 2014). Furthermore, CSNK1A1 has been reported as a mediator of drug resistance to EGFR inhibitors in lung cancer (Lantermann et al., 2015). Unfortunately, potent and selective CSNK1A1 small molecule inhibitors have yet to be reported.

As *CSNK1A1* was among the 3,799 genes subjected to shRNA-mediated knock-down, we used the L1000 data to generate a signature of *CSNK1A1* loss of function. We then queried all compounds in the database against this signature to identify perturbations that phenocopied *CSNK1A1* loss. One unannotated small molecule, BRD-1868, not previously suspected to be a kinase inhibitor, showed strong connectivity to *CSNK1A1* knockdown in two cell types. This suggested that BRD-1868 might function as a novel CSNK1A1 inhibitor. To test this hypothesis, we subjected the compound to kinase specificity profiling, testing its ability to bind to 456 kinases using the Kinomescan assay. Remarkably, the assay confirmed BRD-1868’s ability to bind CSNK1A1 with high specificity and modest potency (K_D_ 2.2 uM). Follow-up enzymatic assays confirmed that BRD-1868 not only binds CSNK1A1, but inhibits its enzymatic activity (Figure 6B). While not yet highly potent, BRD-1868 represents, to our knowledge, the most selective CSNK1A1 inhibitor reported to date, making it an ideal candidate for further chemical optimization. Most importantly, the result highlights the power of using the L1000 Connectivity Map as a starting point for drug discovery – even in the absence of prior examples of the drug class.

### Using L1000 data to assess allele function

The preceding analyses have focused primarily on using the L1000 CMap to functionally annotate small molecule compounds. We next asked whether a similar strategy could be used to annotate the function of an allelic series of genes. Building on results we recently reported (Berger et al., 2016), we sought to determine whether the CMap could distinguish the downstream consequences of overexpression of cDNAs harboring particular somatic mutations observed in human tumors. For example, the ubiquitin ligase FBXW7 is a well-known negative regulator of MYC protein expression. As expected, Connectivity Map analysis found that overexpression of wild-type *FBXW7* strongly connected to knock-down of MYC. In addition, overexpression of 6 mutant alleles found in cancer patients (I347M, V464E, R465C, R465H, A502V, and R505C) all lost this connection to *MYC* loss-of-function, whereas 4 other alleles retained connectivity to *MYC* knock-down (Figure 7A, lower panel). Examination of the substrate-bound FBXW7 crystal structure (Hao et al., 2007) indicated that the mutations predicted by the Connectivity Map to be damaging map to amino acid residues in the FBXW7 substrate-recognition pocket, whereas the non-damaging alleles do not (Figure 7A, upper panel).

**Figure 7.**
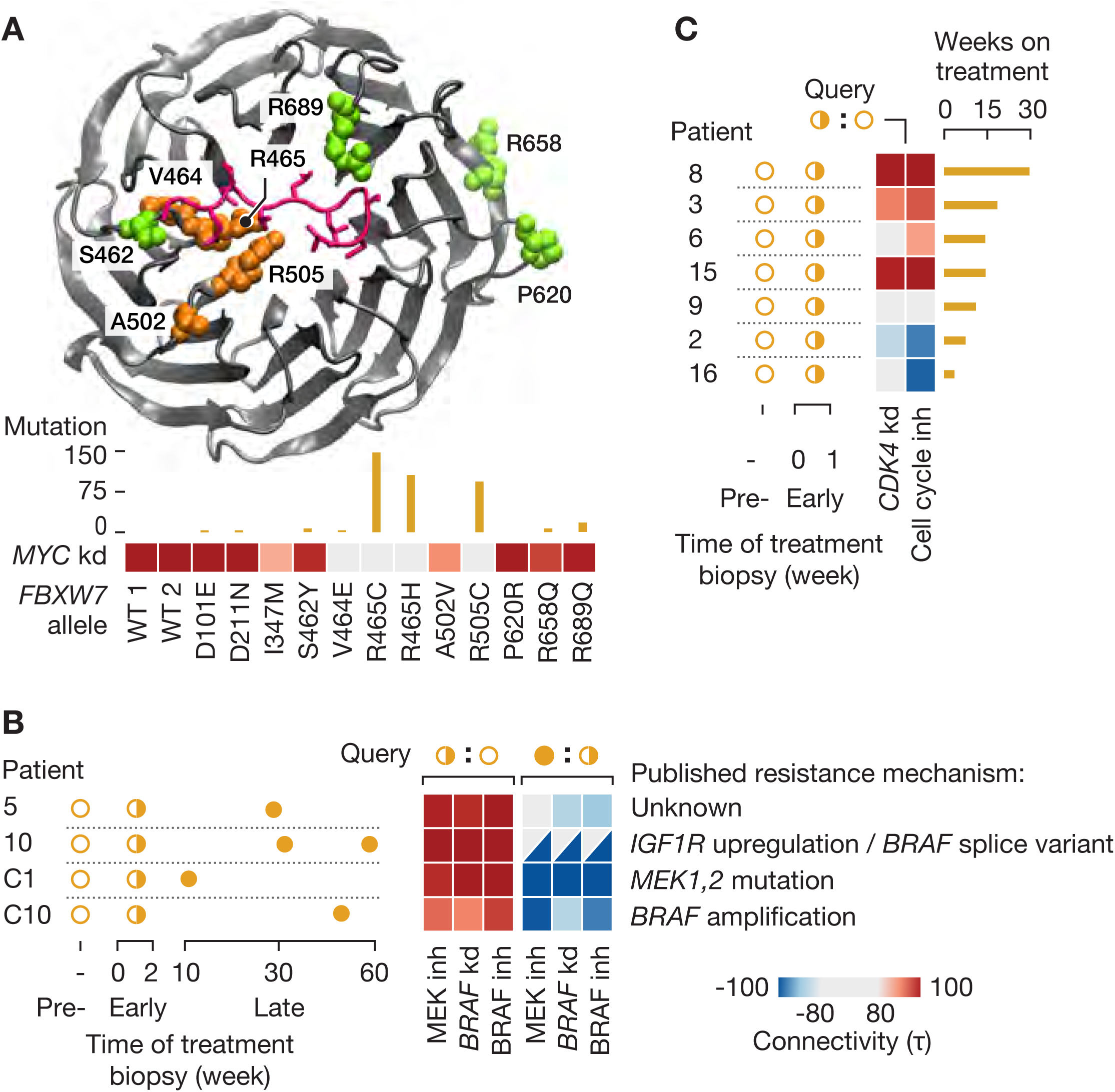
Assessing impact of allelic variants and drug response in clinical trials. **A. Predicting LOF alleles.** A series of clinically-observed mutant FBXW7 alleles were overexpressed in multiple cells types and L1000 profiles were obtained. Overexpression of wild-type FBXW7 connects strongly to MYC shRNA, which is a known target of this ubiquitin ligase. FBXW7 crystal structure was obtained from the Protein Data Bank (2OVQ). Mutations at peptides adjacent to the substrate recognition site (depicted in orange) lose the MYC shRNA connection as shown in the heat map as median tau score across multiple cell types. Bar plot indicates the incidence of each specific mutation in the COSMIC database. **B. Predicting therapeutic efficacy.** Transcriptional profiles of pre-treatment and early on-treatment tumor biopsies were obtained from a published clinical trial of the CDK inhibitor PHA-793887. Differential expression signatures between the two time points were generated for each patient by calculating the log fold change in each gene. CMap query results demonstrated a range of connectivities to negative regulators of the cell cycle. Patients with strong positive connectivity to cell cycle inhibition signatures remained on the clinical trial for a mean duration of 5.25 months, while patients profiles with negative negative correlations were on trial for a mean duration of only two months. **C. Interpreting drug resistance.** Transcriptional profiles of pre-treatment, early on-treatment, and relapse tumor biopsies were obtained from two published melanoma clinical trials of BRAF and MEK inhibitors. Queries from early on-treatment versus pre-treatment biopsies exhibited connectivity to pharmacologic inhibition of BRAF or MEK as well as BRAF shRNA in A375 cells, reflecting target engagement in vivo. MAP kinase signaling was re-activated (as indicated by a strong negative connection to the same CMap reference signatures) in the subset of relapse biopsies with known MAP kinase pathway-related resistance mutations.

The CMap similarly predicted the functional impact of the tumor suppressor KEAP1. Nineteen alleles of *KEAP1* were subjected to L1000 profiling. Whereas wild-type *KEAP1* showed the expected CMap connection to knock-down of its known transcriptional target *NFE2L2*, multiple alleles of *KEAP1* lacked the *NFE2L2* connection, suggesting that these were *KEAP1* loss-of-function alleles. Indeed, a subset of these alleles were recently functionally characterized and reported to result in loss of *KEAP1* function, as predicted by the Connectivity Map analysis (Hast et al., 2014) (Figure S3C, left panel). A similar phenomenon was observed with alleles of the phosphatase PTEN, which negatively regulates PI3K activity. Whereas overexpression of wild-type *PTEN* showed connectivity to signatures of PI3K inhibitors, such connectivity was lost with *PTEN* mutations at residues M35 (mutated in Cowden’s syndrome), G127 (important for active site conformation) and G129 (required for phosphatase activity) (Han et al., 2000; Olschwang et al., 1998) (Figure S3C, right panel). Taken together, these results suggest the feasibility of using L1000 gene expression profiling as a generic read-out of allele function.

### Using CMap to interpret clinical trial results

The CMap resource has been developed to support research, not routine clinical care. However, we hypothesized that there might be potential for the Connectivity Map to inform clinical investigation. Too often, drugs are brought to clinical development without a full understanding of their biological activities in preclinical models, and even more commonly without an understanding of their activity in patients. We analyzed two oncology clinical trials in which tumor samples were obtained before and after treatment, and the resulting gene expression data made publicly available.

In the first study, 21 patients with melanoma were treated with the RAF inhibitors dabrafenib or vemurafenib and 9 patients were treated with dabrafenib plus the MEK inhibitor trametinib (Carlino et al., 2013; Long et al., 2014). Biopsies were obtained prior to treatment and at the time of relapse, and in four patients, early on-treatment biopsies were also obtained. The authors performed expression profiling on the Illumina beadchip platform, and we used those data (GSE50509, GSE61992) as queries to the L1000 Connectivity Map. Comparing the four available early on-treatment biopsies to the pre-treatment biopsies, we observed strong *positive* connectivity to multiple reference readouts of MAP kinase inhibition, consistent with drug-induced silencing of the MAP kinase pathway shortly after treatment. Analysis of patient samples at the time of relapse showed that several patients showed strong *negative* connectivity to these same CMap perturbations, suggestive of reactivation of the MAP kinase pathway – a known mechanism of drug resistance in melanoma (Wagle et al., 2014). One of those patients (patient 10) was indeed shown to have a MAP kinase-activating *BRAF* splice variant, consistent with the Connectivity Map results (Figure 7B). Pathway reactivation was also detected in a resistant tumor with *MAP2K1* mutation (patient C1) and in a resistant tumor with *BRAF* amplification (patient C10).

In the second study, patients with solid tumors were treated with the pan-CDK inhibitor PHA-793887 in a phase I clinical trial. Seven patients from that trial were subjected to gene expression profiling of biopsies obtained pre-treatment and on-treatment using an Agilent microarray platform (Locatelli et al., 2010; Massard et al., 2011). For each patient, the on-treatment expression profile was compared to their pre-treatment profile and the difference used as a signature to query the the Connectivity Map database. The Connectivity Map analysis showed an association between duration of therapy (a proxy for clinical benefit) and connectivity to the overexpression of key negative regulators of the cell cycle such as *CDKN1A* and *CDKN2A*. Strong connectivity was also observed to knock-down of the cyclin-dependent kinase CDK4 – one of the targets of the drug (Figure 7C). Interestingly, the patients with rapidly progressive disease showed *anti-correlation* to this cell cycle inhibition signature, possibly reflective of a feedback mechanism to reactivate the cell cycle in the face of CDK4 inhibition. These results, while reflecting only a small number of patients, are encouraging from a number of perspectives. First, they suggest that while the drug PHA-793887 may be a pan-CDK inhibitor, inhibition of CDK4 may be the most clinically relevant. Such information could be used to guide the development of new chemical analogs. Second, the results suggest that on-treatment biopsy coupled to Connectivity Map analysis may prove useful as an early, molecular readout of target engagement in patients. Such a strategy, if implemented broadly, could bring rich new detail to clinical trials.

### Accessing Connectivity Map data

All of the L1000 Connectivity Map data described in this report are available without restriction to the research community including individuals from commercial entities. To enhance the accessibility and utility of this resource, we have developed a number of computational-visualization tools that enable users to interact with data at multiple levels (from raw data to processed to normalized data), using a variety of methods optimized for technical and non-technical users (e.g., restful Application Programming Interfaces (APIs) for computational biologists and software engineers, and web applications for biologists). The most efficient method of accessing the data and tools is via the secure, cloud-based computing environment that we termed CLUE (**C**onnectivity Map **L**inked **U**ser **E**nvironment), available at https://clue.io. To enable computational researchers to reproduce our findings exactly, code is available at GitHub, and the entire preprocessing workflow is available as a container in the AWS Docker registry. Raw data are also available for download from GEO (accession GSE92742), but users will find it more efficient to interact with the data in the CLUE computing environment.

## DISCUSSION

The study reported here demonstrates the feasibility of a large-scale compendium of functional perturbations of human cells, coupled to a complex, information-rich gene expression read-out. The notion of such a compendium was first proposed in yeast by Friend and colleagues (Hughes et al., 2000), and our pilot Connectivity Map showed that the concept could be extended to mammalian cells (Lamb, 2006). But to be truly useful as a community resource, such a map of molecular connections between cellular states must be comprehensive: if only a limited number of perturbations are profiled, only a limited number of new molecular connections are likely to be discovered.

Our development of the L1000 platform made it possible to scale up the pilot Connectivity Map concept by more than 1,000-fold. By making expression profiling inexpensive, scale up became tractable. The L1000 platform has certain attributes and limitations that are worth considering – particularly when considering transcriptome-wide RNA sequencing as the alternative. Because L1000 is hybridization-based, it is possible to monitor the expression of non-abundant transcripts. While such rare transcripts (e.g., encoding transcription factors) can also be detected by RNA-seq, very high depth of sequencing coverage is needed, and this can become cost-prohibitive. Nevertheless, as sequencing costs continue to drop, RNA-sequencing-based approaches such as Perturb-Seq (Dixit et al., 2016) should be considered for efficient approaches to pooled genetic perturbations in future iterations of the Connectivity Map. We note, however, that L1000 has the advantage of being low-cost even at low-scale, whereas the low cost of RNA-seq can only be achieved at high-scale.

We chose the ∼1000 landmark transcripts in an unbiased manner, based on their orthogonal expression patterns across the diversity of publicly available gene expression data generated by labs across the world, and diverse cell types and disease states they represent. Even so, it is possible that a second generation of L1000 could further improve on the informativeness of the method. Indeed, alternative probe-selection methods have been proposed (Donner et al., 2012). Whether alternative sets of 1,000 transcripts would improve on the ability to discover connections (the primary goal of the CMap) remains to be established.

The ability to infer the expression of genes not directly measured in the L1000 assay was also explored. We found that a simple ordinary least squares model was able to predict the expression of 81% of non-measured transcripts. We recognize that our inference algorithm can likely be improved. Indeed, a recent contest (to be reported in detail elsewhere; data at GEO GSE92743) showed that different computational approaches can significantly improve inference accuracy. An interesting approach would be to develop a cell-type specific inference model, but the extent to which such customized inference improves the ability to discover connections remains to be determined. We also note that while our inference method was successful in the cell types tested, it is conceivable that it could perform less well in cell types highly dissimilar to those used to train the model. As increasing amounts of RNA-seq data are being generated by the community, it will be interesting to see whether such data can be used to further improve L1000 inference.

The 1,319,138 L1000 profiles reported here (yielding 473,647 signatures after combining replicates) represent 42,553 genetic and small molecule perturbations profiled across a variable number of cell types. To our knowledge, this far exceeds any other publicly available resource of cellular perturbation. An important question, however, is the extent to which this large CMap resource can be used to discover important biological connections (e.g., to inform MOA of compounds, to discover pathway membership of gene products, or to connect disease states to pathways and small molecules). In the absence of a true “gold standard,” we attempted to estimate the success rate (and false positive rate) of the CMap approach using a number of metrics.

For example, we used the annotation of protein targets of small molecule drugs and tool compounds to determine whether such targets could be recovered from the CMap. Our results showed that up to 63% of small molecule mechanisms of action could be recovered. The failure to recover the remaining 37% may be explained by a number of factors including ambiguities in annotation of compounds, incomplete target engagement by compounds and genetic reagents, limitations of the L1000 platform, or the inability of transcriptional profiling to recover certain types of connections. Further work should focus on algorithmic improvements that increase the success in recovering known connections. However, even at 63% success, the CMap represents a powerful strategy to discover mechanism of action of new, unannotated compounds. In particular, researchers are often reluctant to perform cell-based small molecule screens because of a concern that discovering MOA may be difficult if not impossible. Our results suggest that Connectivity Map analysis, while not definitive, may rapidly yield MOA hypotheses that can then be experimentally validated.

Indeed, our follow up of several un-annotated compounds indicated that CMap predictions were correct. We focused on kinases simply because of the availability of readily available methods for confirming kinase inhibitory activity of compounds, but there is no reason to believe that the success of the approach is kinase-restricted. Specifically, we found that compounds predicted to be inhibitors of PI3kinase, MTOR, GSK3 or ROCK kinases were in fact inhibitors of those kinases. Perhaps most interestingly, we discovered a highly selective inhibitor of the casein kinase CSNK1A1 – a newly emerging protein essential for survival of certain myeloid malignancies and also implicated in EGFR inhibitor resistance (Lantermann et al., 2015). The compound, BRD-1868, was discovered entirely through computational analysis; no laboratory experiments were needed to generate the CSNK1A1 inhibitory hypothesis. We note that this discovery underscores the value of having a large-scale compendium of genetic and pharmacologic perturbations. Had knock-down of *CSNK1A1* not been profiled in the CMap, and had BRD-1868 similarly not been profiled, the connection between the two would not have been made. Discoveries such as this provide strong rationale for further expansion of the CMap resource.

The discovery of shRNA-small molecule connections also highlights the potential of shRNA-mediated loss-of-function perturbations. Our large-scale analysis of 18,493 shRNA profiles showed that the off-target effects of shRNAs far exceed their on-target effects, consistent with recent reports (Tsherniak et al.). However, the generation of a Consensus Gene Signature (CGS) that identifies gene expression changes common to multiple shRNAs targeting the same gene substantially improved the ability to discover on-target connections by minimizing effects of the off-target component of individual shRNAs. Nevertheless, the CGS procedure is imperfect, and some off-target effects likely remain. Indeed, preliminary studies of CRISPR/Cas9-mediated gene knock-out suggest that genome editing approaches may recover some genetic connections to small molecules that were missed by RNA interference-based perturbation.

Two caveats bear mentioning. First, we and others have recently shown that CRISPR/Cas9-based genome editing results in non-specific toxicity that is directly proportional to the number of cuts to the genome (Aguirre et al., 2016). The extent to which such non-specific effects can be computationally corrected in the context of CMap analysis remains to be determined. This is particularly relevant when performing genetic perturbations in cancer cell lines that often harbor copy number alterations. Second, it remains to be determined whether complete gene knock-out (via CRISPR) or partial knock-down (via shRNA) better phenocopies the effect of a small molecule. For example, complete loss of function may lead to loss of viability, whereas partial loss of function may be tolerated, and thus yield a high-quality transcriptional signature. For the time being, it will therefore likely be wise for further expansion of the CMap to include both shRNA-and CRISPR-mediated perturbational profiles.

Importantly, however, we recognize that genetic loss-of-function (whether by CRISPR or shRNA) and small molecule loss-of-function may not be synonymous. Small molecules often inhibit specific aspects of protein function (e.g., enzyme activity), whereas genetic perturbation generally also results in loss of function entirely – including scaffolding functions that may mediate important protein-protein complex formation. Such scaffolding function may also explain, at least in part, why certain expected connections between small molecules and genetic perturbation of their direct targets were not recovered.

Our analysis of L1000 perturbations across multiple cell types revealed, perhaps not surprisingly, that some perturbations yield universal signatures across cell types, whereas others yield highly cell-type selective gene expression signatures. 43% of compounds active in multiple cell types showed a diversity of cellular responses, arguing for the importance of further expansion of the Connectivity Map into new cell types. The fact that many compounds yield a universal signature regardless of cell type also has important implications. Specifically, the value of continuing to profile such compounds across a large number of cell lines is probably low. In contrast, many yielded unique signatures in neural lineage cells compared to epithelial cancer lines, indicating that an expanded Connectivity Map should continue to explore a diversity of cell lineages and states (e.g., cancer vs. normal). Future iterations of the Connectivity Map might therefore benefit from an adaptive experimental design whereby the selection of future cell lines is chosen based on the performance of an initial set. Furthermore, it is conceivable that computational selection of cell lines most likely to yield new information compared to those already profiled could optimize further growth of the CMap.

Importantly, the Connectivity Map concept is not restricted to mRNA expression as the readout of cellular perturbation. Indeed, the L1000 data reported here were generated as part of the NIH’s Library of Integrated Network-based Cellular Signatures (LINCS) program. Other groups are generating proteomic and other readouts following perturbation. Indeed, recent reports suggest that high-content imaging can also serve as a readout of cellular perturbation (Rohban et al., 2017). Early work by the ChemBank initiative similarly demonstrated feasibility of annotating compounds based on their cellular consequences (Seiler et al., 2008). An important direction for the future will be the integration of such data in hopes of bringing multi-omic readouts to bear on deciphering the functional consequences of experimental perturbations.

Biomedical research in the 21^st^ century reflects a dramatic increase in the sheer amount of data available for analysis, and a commensurate need for increasingly sophisticated computational tools. In the past, researchers would often download genomic datasets to their own computers, and run computational analyses locally. In the era of big data, however, such data downloads become impractical. To facilitate more efficient analysis of CMap data, we have created a cloud-based data storage and analysis system called CLUE, accessible at clue.io. In the CLUE environment, users can access all publicly available CMap data, append any private data, and access a collection of user-friendly analysis apps designed for intuitive use by experimental biologists. Computational biologists can access data using data APIs at clue.io/api. We note that additional L1000 analytical tools developed by others are available through the LINCS data coordinating center website at lincsproject.org.

The Connectivity Map described here represents a dramatic increase in the scale of publicly accessible functional genomic data. Nevertheless, the effort is just a start. A truly comprehensive CMap would expand in multiple dimensions. First, the number of small molecules profiled would increase to include much larger collections (e.g., all FDA-approved drugs and drugs in clinical development, and unoptimized compounds in common screening libraries). Second, the genetic perturbations would include allelic series of important disease-associated genes. The preliminary experiments described here establish the feasibility of creating comprehensive look-up tables of all possible alleles of disease genes, thereby generating functional readouts of genetic variation. Third, while our analyses show that a substantial proportion of perturbations yield universal gene expression signatures, the importance of cellular context is obvious. Accordingly, future iterations of the CMap should explore new cell types including patient-derived iPS cells and genome-edited isogenic cell lines. Fourth, future expansion should include different types of perturbational read-outs. Increasingly, the decreasing cost and increasing throughput of other read-outs (e.g., high content imaging, limited proteomic profiling) make comprehensive Connectivity Map-style perturbational profiling conceivable. An important goal for the years ahead should be to establish which of these alternative data types are most complementary to transcriptional profiling.

Last, we emphasize that the analytical approaches described here represent only an initial approach. Improvements in all aspects of analysis are highly likely to be possible. It is our hope that by making the 1.3 million L1000 profiles freely accessible, the computational biology community will generate new analysis and visualization tools and similarly contribute them to the CMap/LINCS ecosystem.

As with all large-scale community resources, the full potential of the Connectivity Map will only be realized with time. Whether it proves ultimately most useful for elucidating small molecule mechanism of action, for providing functional readouts of allelic series, or for generating new therapeutic hypotheses based on modulation of disease signatures remains to be seen. Such emerging utility should guide the further expansion of a future CMap. However, it seems likely that the explosion of insights into the biological basis of disease, coupled to systematic functional genomic resources such as the Connectivity Map hold potential for a flood of new biologic and therapeutic hypotheses.

## AUTHOR CONTRIBUTIONS

Conceptualization, A.S., R.N., S.M.C., D.D.P., T.E.N., J.L., and T.R.G.; Methodology, A.S., R.N., S.M.C., D.D.P., T.E.N., D.W., I.S., L.H., N.S.G., W.R., J.G.D., and T.R.G.; Software, R.N., T.E.N., J.G., J.K.A., D.L.L., M.K., C.T., and C.F.; Validation, R.N., S.M.C., D.D.P., T.E.N., X.L., and J.F.D.; Formal analysis, A.S., R.N., S.M.C., T.E.N., J.G., D.W., I.S., M.O., O.M.E., C.F., L.H., and J.A.B.; Investigation, S.M.C., D.D.P., X.L., J.F.D., M.D., B.J., D.N., M.B., F.P., A.H.B., D.D, X.W., W.Z., W.R., and X.W.; Resources, F.P., A.H.B., A.S., A.N.B., A.V., D.T., K.A., N.S.G., P.A.C., S.S., X.W., W.Z., S.J.H., L.V.R., J.S.B., S.L.S., J.A.B., and D.E.R.; Data Curation, R.N., S.M.C., T.E.N., J.E.H., Z.L., A.L., and J.A.B.; Writing - Original draft, A.S., R.N., S.M.C., D.D.P., T.E.N., J.E.H., J.A.B., B.W., and T.R.G.; Writing - Review & Editing, A.S., R.N., S.M.C., T.E.N., J.E.H., A.H.B., A.S., D.T., J.L., P.A.C., X.W., S.J.H., L.V.R., J.G.D., J.A.B., D.E.R., B.W., and T.R.G.; Visualization, S.M.C., A.A.T., M.K., and B.W.; Supervision, A.S., D.D.P., J.G.D., B.W., and T.R.G.; Project Administration, A.S., J.R., and T.R.G.; Funding Acquisition, A.S., and T.R.G.

## ACKNOWLEDGMENTS

We thank the Broad Compound Management team, Broad Genetic Perturbation Platform, Harvard Medical School LINCS Center, and LINCS collaborators. We also thank P. Tamayo, J. Bradner and G. Shapiro for helpful scientific discussions. We thank Luminex Corporation for support with the FlexMap 3D system, and Qiagen for assistance with TurboCapture kits. This work was supported in part by the NIH Common Fund’s Library of Integrated Network-based Cellular Signatures (LINCS) program 5U54HG006093 (T.R.G. and A.S.), U54HG008699 (T.R.G. and A.S.) and the NIH BD2K Program grant 5U01HG008699 (T.R.G. and A.S). Additional support was provided by the Howard Hughes Medical Institute (T.R.G.), NIH training grant T32 CA009172 (S.M.C.), KL2/Catalyst Medical Research Investigator Training award from Harvard Catalyst/The Harvard Clinical and Translational Science Center (National Center for Research Resources and the National Center for Advancing Translational Sciences, National Institutes of Health) Award KL2 TR001100 (S.M.C.), and the Conquer Cancer Foundation of ASCO Young Investigator Award (S.M.C.). J.L and W.R.B are shareholders and employees of Genometry, Inc. A.S, R.N, D.D.P, and X.L are shareholders of Genometry, Inc.

## SUPPLEMENTAL LEGENDS

**Figure S1.**
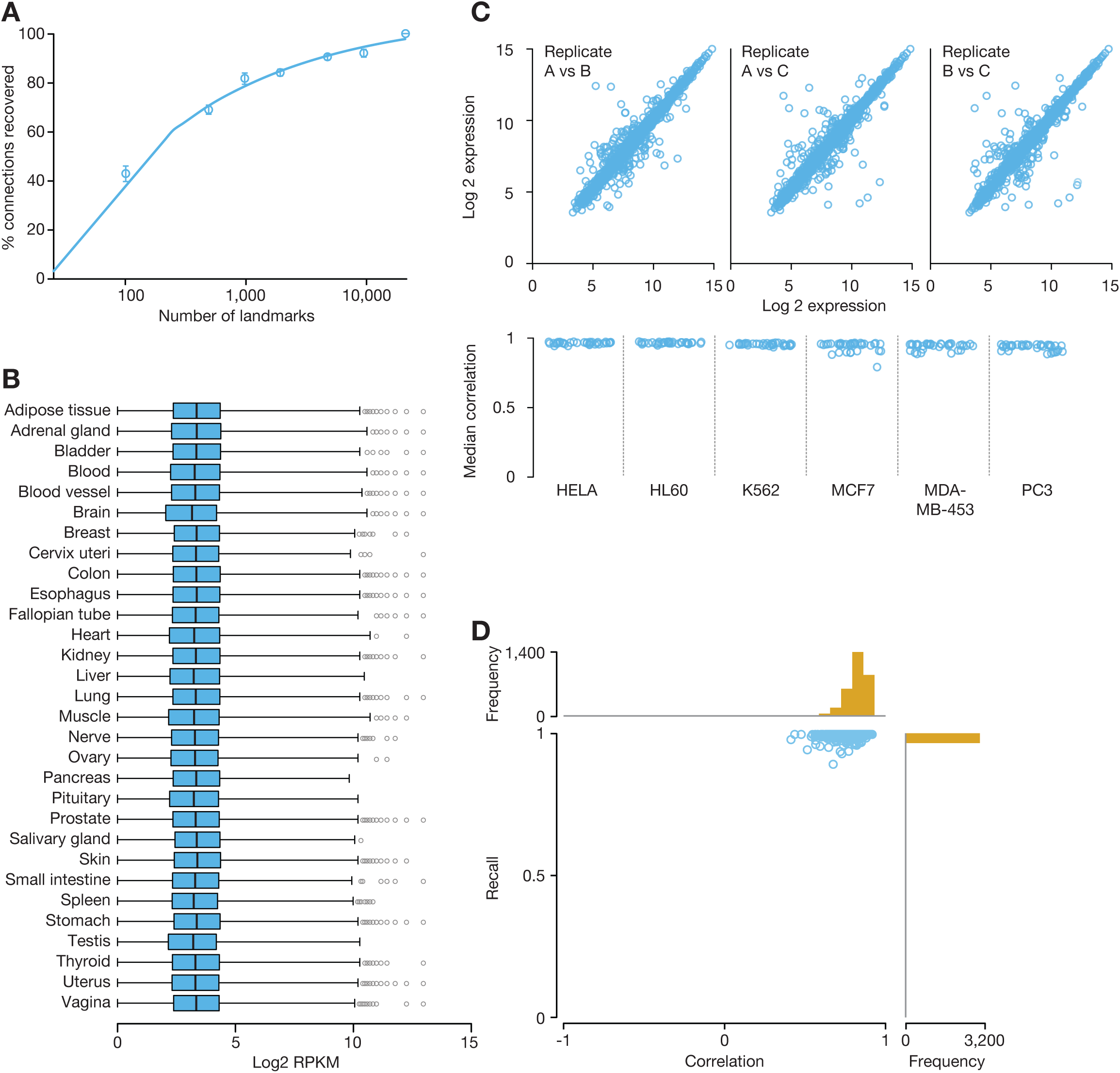
Properties and technical validation of Landmark genes. **A. Evaluating dimensionality reduction**. Simulation showing the mean percentage of 33 benchmark connections recovered from an imputed CMap pilot dataset (+/-SEM) as a function of number of landmarks used in the imputation, indicating that around 1000 landmarks were sufficient to recover 82% of expected connections **B. Landmark gene expression.** Distributions of baseline expression of landmark genes across the 30 tissue types present in dataset DS_GTEx-RNA-seq_. The majority of landmark genes are expressed in each tissue and the distributions of expression are similar across tissues, suggesting that landmarks are not over-optimized to a particular context. **C. Technical replicate reproducibility**. Top: Scatter plots of 3 technical replicates for a sample of commercial MCF7 RNA. Bottom: Distributions of median spearman correlations between technical replicates of commercial RNA samples from 6 cell lines (36 replicates per cell line). Nearly all correlations exceed 0.90, suggesting low sample-to-sample variability. **D. Comparison with RNA-seq**. Scatter plot of replicate recall (RR) vs. cross-platform spearman correlation for 3,176 patient-derived samples profiled on RNA-seq and L1000. 3,103/3,176 samples (98%) had a RR < 0.01 (indicating 99th percentile) and all but 5 samples (99.84%) had a RR < 0.05, suggesting high similarity between the two platforms.

**Figure S2.**
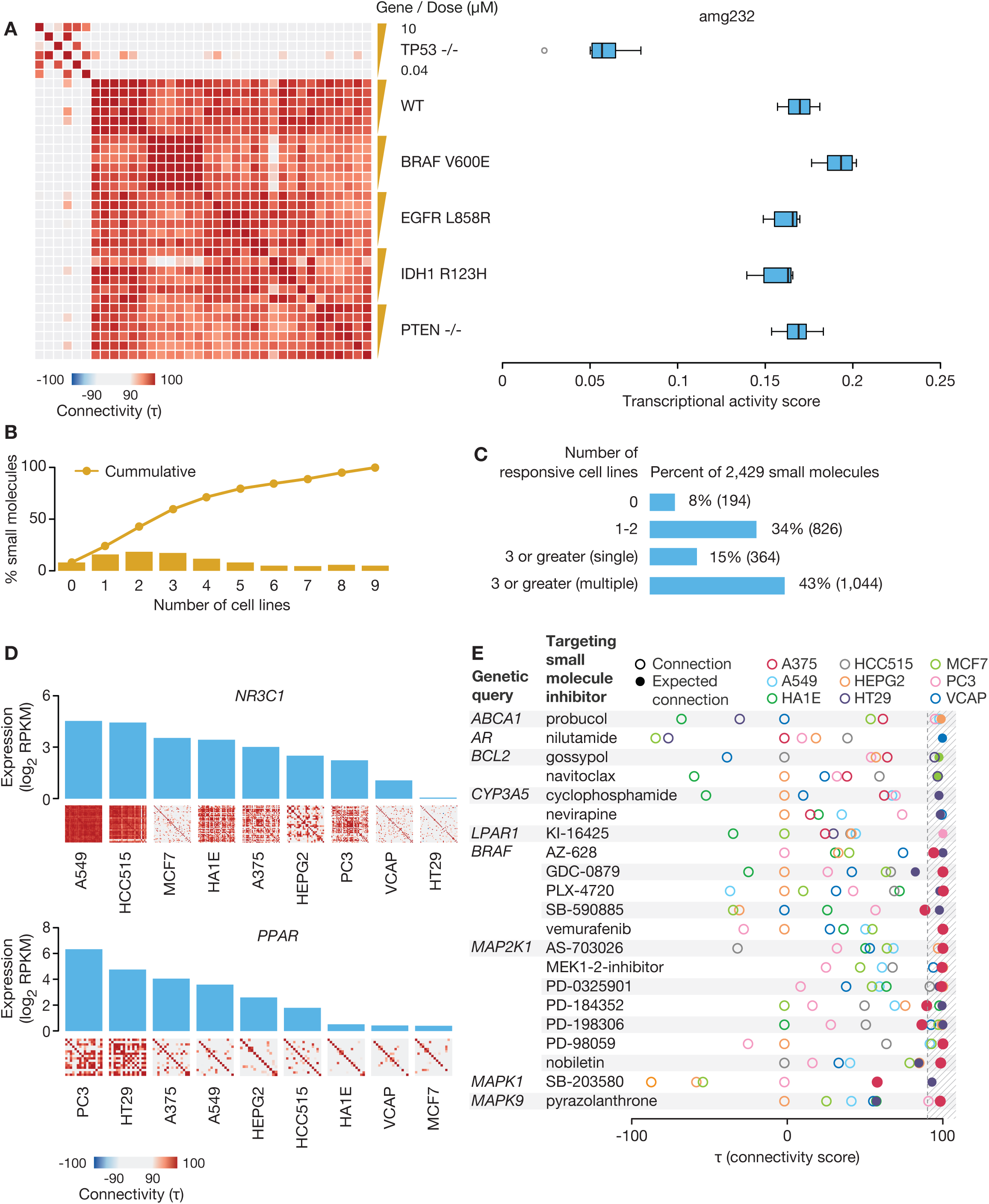
Characteristics of perturbational signatures. **A. MDM2 inhibitor AMG-232 shows TP53-dependent transcriptional response**. Left: Correlation matrix of AMG-232’s signatures across cell lines at 6-point dose ranging [0.04-10 uM]. Signatures in background of *TP53* homozygous deletion lose correlation with signatures in *TP53* wild type background. Right: Boxplots of AMG-232 TAS. TAS is notably lower in background of *TP53* homozygous deletion compared to wild type. **B. Cell context and transcriptional activity.** The bars show the percentage of 2,429 small molecule compounds that produced high TAS signatures in a specified number of cell lines (TAS ≥ 0.2). The cumulative percentages are shown in blue. 92% of compounds produce a high-TAS signature in at least one cell line. **C. Multiplicity of signatures**. The bars indicate the percentage of 2,429 compounds that gave a single or multiple signatures across all the cell lines in which each compound was active (minimum of 3 cell lines). Compounds that were active in fewer than three cell lines are captured in the first two bars. Only 15% of compounds produced a single signature, suggesting that transcriptional response is dependent on cell context for a majority of compounds. **D. Context-specific interconnectivity**. The bar plot shows the baseline expression of the gene *NR3C1*, the target of these compounds, across the 9 core cell lines. The small heat map below each bar shows the interconnectivity of the 44 members of the glucocorticoid agonist PCL in the same cell line. The interconnectivity is highest in the cell lines in which the target is most highly expressed. Right) The bar plot shows baseline expression of the gene *PPARG*, which encodes the target of PPARG agonists. The thumbnail heatmaps show the inter-connectivity of 16 members of the PPARG receptor agonist PCL. **E. Context-dependent connection between compounds and their gene targets.** The raster plot shows the connectivity between the indicated gene knockdown and its targeting compounds, where each circle is a cell line. Filled circles indicate cell lines in which the compound-gene connection is expected, either due to mutation in (*BRAF*, MAPK genes) or high expression of (all others) the target gene. The strong connections are enriched for the expected cell lines, suggesting that CMap connectivities can be indicative of context specificity.

**Figure S3.**
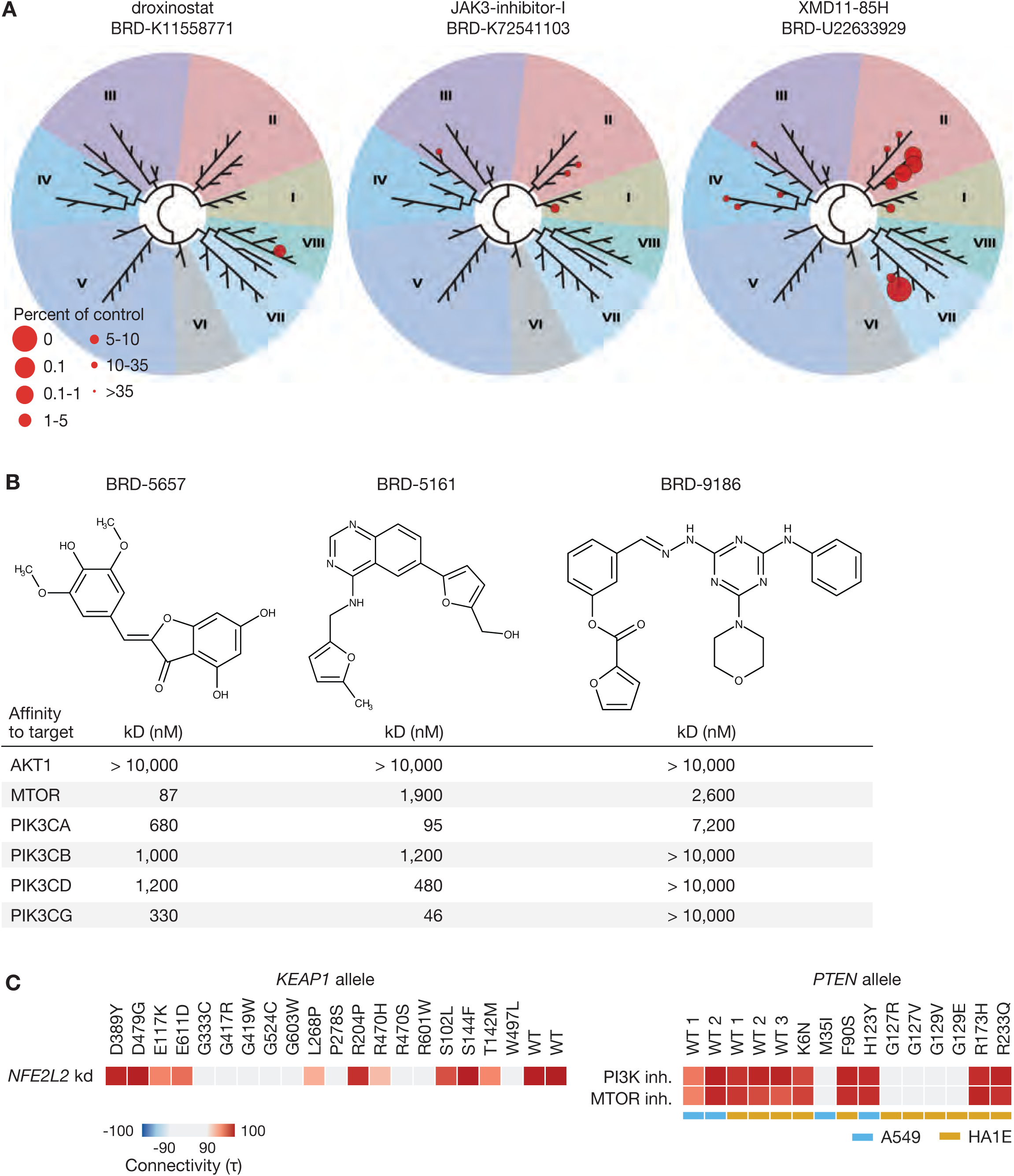
Experimental follow-up of CMap hypotheses and functional annotation of allelic variants. **A. Confirmation of predicted bromodomain inhibitors.** TREEspot selectivity profile indicating the inhibitory activity of droxinostat, XMD11-85H, and BRD-1103 against the 32 bromodomains tested by the BROMOmax assay. All three compounds were active against at least one bromodomain. Images generated using TREEspot Software Tool and reprinted with permission from KINOMEscan, a division of DiscoveRx Corporation. **B. Confirmation of predicted PI3K/MTOR inhibitors**. Chemical structures and affinities (kD) between three predicted PI3K/MTOR inhibitors and *AKT1*, *MTOR*, and 4 *PI3K* isoforms, as measured by the Kinomescan assay. All three compounds show at least micromolar affinity for a subset of the predicted targets, with two compounds showing nanomolar affinities. **C. CMap connectivities correspond to loss of function mutations**. Left: Heat map showing the connectivity between the over-expression signatures of various *KEAP1* alleles in A549 cells to the knockdown of *NFE2L2* in A549 cells. Right: Heat map showing the connectivity between the over-expression signatures of various *PTEN* alleles and the PI3K and MTOR inhibitor classes. For both *KEAP1* and *PTEN*, mutations at known active residues important for protein function result in loss of expected connectivity.

## SUPPLEMENTARY INFORMATION

### 1. QUICK GUIDE TO WHERE TO FIND METHODS IN THIS SUPPLEMENTAL TEXT

1. **Reduced representation of transcriptome. “***We therefore used those 25 signatures to query the imputed DS*_*CMAP-AFFX*_*dataset for each value of k, counting how often we recovered the connections observed in the original dataset at a comparable rank based on the Kolmogorov-Smirnov statistic”:* ***Methods section 3***
2. **L1000 assay platform.**

a. *“Thus, each bead was analyzed both for its color (denoting landmark identity) and fluorescence intensity of the phycoerythrin signal (denoting landmark abundance). Because only 500 bead colors are commercially available, we devised a strategy that allows two transcripts to be identified by a single bead color”:* **Methods section 4**
b. “T*he final assay, which we call L1000, contains 1,058 probes for 978 landmark transcripts and 80 control transcripts chosen for their invariant expression across cell states”:* **Methods section 4**
3. **Optimization and validation of L1000. “***In designing landmark-specific oligonucleotide probes, we followed several computational procedures that maximized matches to the target DNA sequence while minimizing non-specific hybridization”:* **Methods section 5**
4. **CMap query methodology**

a. *“Each of the signatures in the database represents a weighted average across the 3 biological replicate perturbation”:* **Methods section 9**
b. *“We therefore developed a Connectivity Score”*: **Methods section 9**
5. **Characterizing small-molecule function. “***This led to 58,820 expected relationships that could plausibly be recovered in the CMap-L1000v1 compendium”:* **Methods section 12**
6. **Defining perturbagen classes (PCLs)**

a. *“These perturbagen classes (PCLs) were then further refined by excluding from the PCL any compounds that failed to empirically connect with their cognate class members based on L1000 connectivity analysis”:* **Methods section 13**
b. *“We next asked whether drugs with established MOA also had unexpected, strong, and selective connections to a validated PCL. Selectivity was defined as the fraction of PCLs to which a drug failed to connect at a given τ threshold. 132 drugs (3.9%) had such off-target connections”:* **Methods section 13**
7. **Cellular context**. “*Of 1,399 (58%) compounds active in at least 3 cell lines, 26% (corresponding to 15% of all compounds) produced highly similar signatures across the entire panel, whereas perhaps not surprisingly, the remainder were active in only 1 or 2 cell lines or produced a diversity of cellular signatures”:* **Methods section 14**.
8. **Identifying bioactive subsets of small-molecule screening libraries. “***We found that whereas 2,232/2,429 (92%) established drugs yielded a strong L1000 transcriptional response defined as Transcriptional Activity Score (TAS) >0.2”:* **Methods section 15**

### 2. DATA ACCESS - LIST OF DATASETS USED AND WHERE TO DOWNLOAD THEM

All datasets used or referred to in this manuscript are available for download by all from the NCBI Gene Expression Omnibus (GEO) under accession number GSE92742.

| Name | Brief description / use case |
| --- | --- |
| **Datasets** |  |
| Dataset—DS<sub>GEO</sub> | Collection of publicly available gene expression profiles on Affymetrix HGU133A arrays that was used to select landmark genes and train the inference model.<br>*rows: 22,268 features*<br>*columns: 12,301 samples* |
| Dataset—DS<sub>GEO-OLS</sub> | The matrix of weights learned by training the L1000 inference algorithm, ordinary least squares (OLS) linear regression, on DS<sub>GEO</sub>.<br>*rows: 21,290 inferred features*<br>*columns: 978 landmark genes + intercept = 979* |
| Dataset—DS<sub>CMAP-AFFXB01</sub> | Pilot connectivity Map dataset of 455 gene expression profiles of human cell lines treated with 164 small molecule compounds that were used in simulations to determine the optimal number of landmarks.<br>*rows: 22,268 features*<br>*columns: 455 samples* |
| Dataset—DS<sub>reproducibility</sub> | Samples of purified total RNA from six human cancer cell lines, purchased from Life Technologies, were subjected to L1000 profiling. L1000 expression profiles were generated consisting of 12 technical replicates for each of the six cell lines, all done in three consecutive, independent LMA batches, yielding 36 replicate profiles per cell line and a total of 216 total profiles (6 cell line x 12 replicates x 3 batches). These profiles were used to assess the technical reproducibility of the L1000 assay.<br>*rows: 978 landmark genes*<br>*columns: 216 samples* |
| Dataset—DS<sub>LM-KD</sub> | We evaluated if probes designed against individual target landmark genes worked in the multiplexed gene assay format. To assess the specificity of L1000 landmark probe measurements, we procured shRNAs from The RNAi Consortium (TRC) that targeted landmark genes, treated MCF7 and PC3 cell lines with these shRNAs. The resulting dataset contains, as columns, an individual shRNA targeting a landmark gene performed in either MCF7 or the PC3 cancer cell line. Rows are z-scores of all measured landmark genes. For each gene in each sample, we computed differential expression values (z-scores) by comparing the gene's expression value in the given sample to that same gene's expression values in all other samples.<br>*rows: 978 landmark genes*<br>*columns: 955 samples* |
| Dataset—DS<sub>GTEX-RNA-seq</sub> | Compendium of 8,555 RNA-seq samples obtained from the GTEx consortium (version 6). A subset of 3,176 of these samples were also profiled on L1000 and were used to compare the two platforms.<br>*rows: 12,320 genes*<br>*columns: 8,555 samples* |
| Dataset—DS<sub>GTEX-L1000</sub> | From the GTEx version 6 collection, a subset of 3,176 samples were generously donated by the GTEx consortium for profiling in the L1000 assay.<br>*rows: 970 landmark + 11,350 inferred = 12,320 genes*<br>*columns: 3,176 samples* |
| Dataset—DS<sub>GTEX-RNA-seq-lmonly</sub> | For convenience of use during assay validation and inference testing, we separated out from the overall GTEx RNA-seq dataset (DS<sub>GTEX-RNA-seq</sub>), the subset of landmark genes and refer to it as DS<sub>GTEX-RNA-seq-lmonly</sub> in the methods. Note that the GTEx pre-processing provided mappings for 970 of the landmark genes.<br>*rows: 970 landmark genes*<br>*columns: 8,555 samples* |
| Dataset—DS<sub>GTEX-RNA-seq-INF</sub> | Inferred version of DS<sub>GTEX-RNA-seq</sub> dataset generated by applying DS<sub>GEO-OLS</sub> to DS<sub>GTEX-RNA-seq-lmonly</sub>.<br>*rows: 970 landmark + 11,350 inferred = 12,320 genes*<br>*columns: 8,555 samples* |
| Below are the datasets corresponding to the main CMap-L1000 v1 resource generated and used in this paper. For more details on download and format of datasets deposited in GEO, please see: https://clue.io/GEO-guide |  |
| GSE92742_Broad_LINCS_Level1_LXB_n1403502.tar.gz | **Level 1 - RAW:** Archive of raw LXB files.<br>*1,403,502 scans* |
| GSE92742_Broad_LINCS_Level2_GEX_delta_n49216x978.gctx.gz &<br>GSE92742_Broad_LINCS_Level2_GEX_epsilon_n1269922x978.gctx.gz | **Level 2 - GEX:** Deconvoluted expression profiles. Data are provided in two separate files, one each for two slightly different iterations of the landmark gene set (delta and epsilon). Note that the vast majority of data was generated using epsilon. Very poor or corrupt LXB files were discarded.<br>*rows: 978 landmark genes*<br>*columns: 1,319,138 profiles* |
| GSE92742_Broad_LINCS_Level3_INF_mlr12k_n1319138x12328.gctx.gz | **Level 3 - QNORM:** LISS and quantile-normalized expression profiles.<br>*rows: 978 landmark + 11,350 inferred = 12,328 genes*<br>*columns: 1,319,138 profiles* |
| GSE92742_Broad_LINCS_Level4_ZSPCINF_mlr12k_n1319138x12328.gctx.gz | **Level 4 - ZS:** Differential expression profiles.<br>*rows: 978 landmark + 11,350 inferred = 12,328 genes*<br>*columns: 1,319,138 profiles* |
| GSE92742_Broad_LINCS_Level5_COMPZ.MODZ_n473647x12328.gctx.gz | **Level 5 - MODZ:** Replicate-collapsed differential expression signatures.<br>*rows: 978 landmark + 11,350 inferred = 12,328 genes*<br>*columns: 473,647 signatures* |
| **Laboratory Protocols** |  |
| Doc D01—L1000 assay Standard Operating Protocol<br>PDF | The full protocol for the laboratory execution of the L1000 assay. |
| Doc D02—Cell culture and treatment conditions<br>PDF | Details of culture conditions used for each of the core cell lines in the current dataset. Includes cell plating density and media used. |
| **Code : http://github.com/cmap/cmapM** |  |
| Code—Preprocessing Level 1 (RAW) to Level 2 (GEX) | Deconvolute 978 gene expression values from 500 Luminex bead colors. |
| Code—Preprocessing Level 2 (GEX) to Level 3 (QNORM) | 1. To mitigate non-biological variation, adjust samples according to 80 in-well invariant control genes and rescale such that all samples follow the same empirical distribution.<br>2. Infer the expression of 11,350 non-measured genes from the 978 landmarks. |
| Code—Preprocessing Level 3 (QNORM) to Level 4 (ZS) | Obtain measures of differential expression (z-score) for each gene in each sample. |
| Code—Preprocessing Level 4 (ZSPC) to Level 5 (MODZ) | To mitigate outlier samples and obtain a more robust signature of a perturbation, we collapse level 4 profiles into a composite signature using a weighted averaging procedure. |
| Code—infer_ols | Given a new profile with the 978 landmark gene measurements, use $DS_{GEO-OLS}$ to impute the values of genes |
| Code—train_ols | Given the $DS_{GEO}$ and landmark gene identities, train an OLS model and generate $DS_{GEO-OLS}$ |

### 3. REDUCED REPRESENTATION OF THE TRANSCRIPTOME

#### Dataset for Landmark selection

We assembled a large, diverse collection of 12,063 gene expression samples profiled on Affymetrix HG-U133A microarrays from the Gene Expression Omnibus (GEO) (Edgar R et al., 2002). These data were used to identify the subset of universally informative transcripts to be measured, which we term ‘Landmark Genes’ (Dataset DS_GEO_).

**Simulations to determine appropriate number of landmarks**

We sought to determine the optimal number of landmarks. To address this, we asked what number of landmarks would optimally recover the observed connections seen in the pilot Connectivity Map dataset based on Affymetrix arrays (Dataset DS_CMAP-AFFXB01_). We assembled a collection of 25 query signatures based on prior external work that yielded 33 robust and expected connections in the pilot DS_CMAP-AFFXB01_dataset (Table S1). At each value of k (ranging 100-10,000), we generated an imputed version of the this dataset using OLS regression (trained on samples from DS_GEO_) with a random selection of k landmarks as the independent variables, queried it with the benchmark signatures, and assessed the percentage of connections that were recovered. We repeated this procedure 10 times for each choice of k. Figure S1A shows the mean percentage of connections recovered as a function of k, suggesting that on average 82% of connections are recovered at k=1,000 landmarks.

#### Landmark genes - selection procedure

We adopted a data-driven approach to select 1,000 landmark genes using the DS_GEO_ dataset. Because the dataset contains a non-uniform representation of various aspects of biology (for example certain tumor types such as breast and lung cancer were disproportionately represented), we applied Principal Component Analysis (PCA) as a dimensionality reduction procedure to minimize bias toward any particular lineage or cellular state. In a reduced eigenspace of 386 components (which explained 90% of the variance), cluster analysis was performed on the loadings to identify clusters of commonly co-regulated transcripts. We applied an iterative peel-off procedure to select the centroids (Tseng and Wong 2005). At each iteration we identified the most concordant clusters via k-means. For each cluster the transcript closest to the centroid was selected as a candidate landmark gene. All cluster members were subsequently dropped and the procedure was repeated to identify additional clusters in the remaining feature space.

#### Landmark genes - baseline expression of landmark genes across a diversity of tissue types

Our procedure for selecting Landmark Genes was data-driven and the simulations presented above indicate that both the landmark and inferred genes capture relevant information about cell state. However, given a new state, any inference algorithm will only work if a fair number of the landmark genes are expressed in that state. We examined expression across lineage using the Genotype Tissue Expression (GTEx) RNA-seq dataset (DS_GTEx-RNA-seq_) of 3,176 patient-derived expression profiles from 30 different tissue types (Figure S1B). We quantified the expression levels of the landmark genes reported in the dataset and observed that at a RPKM threshold of 1 at least 86% of Landmark Genes are expressed in each of the 3,176 samples (with an average of 92% expressed in each sample), and that range of expression is similar across tissue types.

#### Landmark genes - functional enrichment analysis of landmark content

Our data-driven procedure suggested genes to include as landmarks based on analysis of the 12,063 sample compendium DS_GEO_. We then asked if genes suggested by this data driven approach were enriched in particular known biological pathways or categories.

For every landmark gene we accessed from the NCBI Entrez database its current gene description and family assignment. We also annotated every landmark gene with the pathway (as defined in MSigDB) in which it is thought to function (when available). Finally, we looked up its biological/molecular category from Gene Ontology (GO). These annotations were analyzed for functional enrichment to ask if the landmarks, when considered as a set, are dominated by a few functions or if on the whole they map to many different functions. For example, at one extreme the transcriptionally active genes could belong to basic regulatory processes (e.g transcription factors).

To do this analysis we intersected the 978 landmarks with a database of gene sets compiled in Gene Ontology using the hypergeometric statistic (gene to GO gene ontology, conditional test for over-representation). We used the R Bioconductor package GOstats (v2.36.0) and the ontology from GO.db (v3.2.2). The results shows that while some categories are enriched (e.g ATP binding, nucleoside/nucleotide activity, transcription factor binding, kinase regulator activity) the percentage of the 978 genes that are in any such set is small. While we did observe a number of classes to be enriched in the landmark genes, these categories tend to be generic (e.g. enzyme binding, protein kinase binding, catalytic activity, ATP binding) and/or contain only a small fraction of the landmark genes (e.g. protein kinase binding, which contains 84 of 978 landmarks). Taken together, we did not find any particular functional category dominating the list of landmarks chosen.

### 4. L1000 ASSAY PLATFORM

#### Overview of the L1000 assay

Having established through simulations that measuring approximately 1,000 landmarks was sufficient to capture the majority of information encoded in genome-wide expression profiles, we next sought to develop a laboratory method capable of actually measuring 1,000 transcripts at low cost. For this purpose, we adapted a method we previously reported involving 90-plex ligation-mediated amplification (LMA) followed by capture of the amplification products on fluorescently-addressed microspheres (beads) (Peck et al., 2006). We extended this method to a 1,000-plex reaction. Briefly, cells growing in 384-well plates were lysed and the mRNA transcripts captured on oligo-dT-coated plates. 1,000 distinct locus-specific oligonucleotides harboring a unique 24-mer barcode sequence were then used to perform an LMA reaction, and the biotinylated LMA products were detected by hybridization to optically addressed polystyrene microspheres (beads), each coupled to an oligonucleotide complementary to a barcode, and staining with streptavidin-phycoerythrin. Each bead was analyzed for its bead color (denoting the landmark identity) and its phycoerythrin fluorescence intensity (denoting the landmark transcript abundance) using a Luminex FlexMap 3D system. Because only 500 unique bead colors are commercially available we developed a procedure called TagDuo that allows two genes to be detected on a single bead color (see below). The final L1000 assay contained probes for 978 landmark transcripts and 80 control transcripts chosen for their invariant expression (see below).

#### Probes and primers

Each transcript of interest was targeted with an upstream and downstream probe pair. Upstream and downstream probes were each designed with a 20nt gene specific region (40nt contiguous sequence per probe pair), a unique identifying barcode, and a universal primer site. The gene specific sequences were blasted against the human genome to verify that each is unique to the targeted gene of interest, as described in the steps below. In addition to gene specific sequence, upstream probes contained a T7 primer site, and a 24-nucleotide (nt) barcode, and downstream probes, which were 5’ phosphorylated, contained the T3 primer site. Barcode sequences are shown in supplementary Table S2. Probes were synthesized by IDT (Integrated DNA Technologies).

We followed an iterative process of probe design followed by empirical probe validation, as follows, until we achieved ∼1,000 landmark genes with a validated probe.

1. Landmark genes proposed based on computational analysis.
2. For each gene, select a 40 base sequence using the following design principles, then split into two 20-mers

a. Empirical probe design rules:

i. the region must be contiguous with no gaps
ii. must be 3’ biased to minimize RNA degradation
iii. choose regions with few repeats to minimize cross-reactivity
3. Perform computational sequence QC by aligning against human reference genome (assembly HG19) using BLAT (Kent 2002)

a. Ensure a perfect alignment to intended gene′s reference sequence
b. Check for non-specific alignment of the probe sequence to other genes
c. If either checks (a) or (b) fail, then redesign the probe sequence
4. Build upstream and downstream probes using T7 and T3 primer sites and FlexMAP tag

a. T7 primer site 5’ TAA TAC GAC TCA CTA TAG GG 3’
b. T3 primer site 5’ TCC CTT TAG TGA GGG TTA AT 3’
c. Uni-bio-T7 5’/5Bio/TAA TAC GAC TCA CTA TAG GG 3’
d. Uni-T3 5’ ATT AAC CCT CAC TAA AGG GA 3’

#### Cell lysate preparation

Cells were cultured in appropriate media and 40 μl was transferred into each well of a 384-well clear bottom, tissue culture treated plate with an automatic liquid handler (for more detail about determining the optimum cell densities, see Doc D02). Plates were incubated at 37°C, 5% CO_2_. Cells are either treated with chemical or genetic perturbations, the details of which are reported in section 6 below. For cell lysis, media was removed from the wells without disturbing the cells and 25 μl/well of TCL Lysis Buffer (Qiagen) was added. Plates were sealed with adherent foil seals and incubated at room temperature for 30 minutes prior to storage at -80°C.

#### Coupling barcodes to Luminex beads

To detect gene-specific sequences, Luminex beads were coupled to DNA barcodes complementary to each barcode used in our collection of probes. Because Luminex produces 500 distinct bead colors and the L1000 set consists of 978 genes, 2 barcodes were coupled to beads of each color (see below); this was done in separate batches -one barcode per batch - and then the pairs were mixed in a 2:1 ratio prior to use. Luminex magnetic beads were added in 500 μl aliquots to each well of 96 deep-well plates. Beads were pelleted and resuspended in 62.5 μl binding buffer (0.1 M 2- [N-morpholino]ethanesulfonic acid; pH 4.5), to which was added 100 pmol capture barcode. 6.25 μl of freshly prepared 10 mg/ml aqueous solution of 1-ethyl-3-(3-dimethylaminopropyl) carbodiimide hydrochloride (Pierce, Milwaukee, WI, USA) was added to each well followed by incubation at room temperature in the dark for 30 minutes. This step was repeated and then 180 μl 0.02% Tween-20 was added. Beads were pelleted and washed in 0.1% SDS in TE, pH 8.0 buffer. Beads were stored in TE in the dark at 4°C for up to one month. Mixtures of beads were freshly prepared in 1.5X TMAC buffer (4.5 mol/l tetramethylammonium chloride, 0.15% N-lauryl sarcosine, 75 mmol/l tris-HCl [pH 8.0], and 6 mmol/l EDTA [pH 8.0]).

#### Ligation-mediated amplification

For mRNA capture, 20 μl lysate was transferred to Turbocapture (Qiagen) plates coated with oligo dT. Following a 60-minute incubation at room temperature, unbound lysate was removed by inverting the plates onto a highly absorbent towel followed by centrifugation at 1000 rpm for one minute. First-strand cDNA was prepared from the mRNA by adding 5 μl master mix consisting of # units M-MLV reverse transcriptase and # umol/l of each dNTP. Plates were incubated at 37°C for 90 minutes. Probes were annealed to the first-strand cDNA using 5ul Probe Anneal master mix, which contains 100 femtomole of each probe in 1X Taq ligase buffer (New England BioLabs, Inc.). Denaturation was accomplished by incubating the plates at 95°C for 2 minutes and then decreasing the temperature from 70°C to 40°C over a 6-hour period. Plates were then inverted onto an absorbent towel and spun at 1,000 RPM for 1 minute to remove unbound probe.

To ligate juxtaposed probe pairs, 5 μl mix containing 2.5 units Taq DNA ligase (New England Biolabs) in ligase buffer was added, plates were sealed, and incubation proceeded at 45°C for 1 hour followed by 65°C for 10 minutes. The plate wells were emptied as described above, and the resulting amplification templates were subject to PCR using T3 and 5’-biotinylated T7 universal primers. PCR was initiated by adding 15 μl master mix, containing 1.5 umole of each primer, 2.4 nmol of each dNTP, and 4.8e-4 units of HotStarTaq in reaction buffer. Plates were sealed and loaded into a Thermo Electron MBS 384 Satellite Thermal Cycler. Initial denaturation was performed at 95°C for 15 minutes, and then the plates were subjected to 29 cycles as follows, one minute per step: 92°C (denature), 60°C (anneal), 72°C (elongation). The resulting amplicons were gene-specific, barcoded, and biotinylated.

#### Hybridization of amplicon to bead

Because a sequence complementary to the barcode on each probe has also been coupled to a Luminex bead, the amplicons (and hence the gene-specific sequence) can be identified by hybridization to the beads. A volume of 5 μl of PCR amplicon was transferred to a well containing 30 μl of L1000 bead mix (about 350 beads/region/well). The plate was sealed and incubated at 95°C for 2 minutes to denature the DNA. Incubation continued at 45°C for 18 hrs. Beads were pelleted, washed, and stained with 20 μl of 10 ng/ul streptavidin R-phycoerythrin conjugate (Molecular Probes) in 1× TMAC buffer (3 mol/l tetramethylammonium chloride, 0.1% N-lauryl sarcosine, 50 mmol/l tris-HCl [pH 8.0], 4 mmol/l EDTA [pH 8.0]) at 45°C for ten minutes.

#### Tag Duo dual detection and peak deconvolution

The Luminex FlexMap 3D platform is capable of detecting 500 different bead colors while the L1000 assay needs to measure ∼1,000 mRNA transcripts. One option would be to read these in 2 different detection sets, each of 500. However, that detection strategy would introduce inevitable batch effects and also reduce throughput of detection by half.

Therefore we devised a strategy that allowed two different transcripts to be identified using a single bead color. The 978 landmarks were divided into pairs and barcodes representing each gene were coupled to beads of the same color (one gene per bead). Genes coupled to the same bead type (color) in two separate batches were combined in a ratio of 2:1 prior to use. When the beads are hybridized with the sample templates and analyzed by the Luminex scanner, two values are obtained from each bead: one indicating the color of the bead and the other indicating the intensity of the signal, which is a reflection of the expression of the gene. Identification of the bead color associates the intensity to the correct gene pair and signal intensity provides a measure of the abundance of the transcripts of the two genes. Deconvolution of the composite fluorescent intensity signal into its component gene expression values is done computationally as described in 8. Data Pre-Processing – Raw Data to Signatures (see also Figure 1B). To make it easier to resolve the peaks, rather than pairing genes at random, during design of the L1000 system, we optimized pairing of genes to maximize the average difference in their expression levels across the training GEO compendium.

#### Detection

Hybridization of amplicon to complimentary barcodes was detected using a Luminex FlexMap 3D flow cytometer, which detects both bead color (i.e., transcript identity) and the biotin label on the probe (i.e. transcript abundance; as measured by the phycoerythrin channel). Analysis was done using a sample volume of 40 μl.

#### Invariant genes as controls for data QC and normalization

We developed a set of internal controls to assess quality, to provide real-time feedback during the scanning process, and to use in normalization. Importantly, rather than using a single “housekeeping” gene (e.g. GAPDH), we adopted an approach that utilizes control values across the entire spectrum of gene expression. We adapted the approach described in the Illumina BeadChip studio (Illumina, n.d.) by defining a set of genes that are rank invariant across all samples. To identify these genes, we analyzed human gene expression profiles from DS_GEO_ and selected genes whose expression is relatively invariant (coefficient of variation < 10%) across a variety of tissue types and experimental conditions. To further minimize the variance, rather than picking single genes as invariants, we grouped the genes into 10 sets of 8 genes each based on their level of expression across all samples. The 10 gene sets were ordered by increasing levels of expression, with the first level corresponding to genes with the lowest expression and the tenth level to genes most highly expressed. Because these gene sets exhibit a consistent expression pattern, they can be used to adjust the data for non-biological variation. Importantly, in addition to being useful for data normalization, the invariant genes provide a simple quality check in real time as detection occurs, which is valuable in a high-throughput process.

## 5. OPTIMIZATION AND VALIDATION OF L1000

The L1000 assay is optimized for the rapid measurement of endogenous gene-expression of selected transcripts. In addition, substantial automation was introduced into the assay protocol so as to enable the massive scale-up in data generated with minimal personnel needs—a laboratory team of ∼4 generated most of the 1 M profiles over a 48 month period.

Given the notably different technological approach to building the compendium, we evaluated the operating characteristics of the platform to establish reproducibility, sensitivity and stability of the platform.

### Reproducibility of L1000—using reference mRNA

Samples of purified total RNA from six human cancer cell lines, purchased from Life Technologies, were subjected to L1000 profiling. L1000 expression profiles were generated for six cell lines in 3 independent LMA batches, each with 12 technical replicates, for a total of 216 total profiles (6 cell line x 12 replicates x 3 batches = 216). Within each cell line, we computed the Spearman correlation between all pairwise combinations of replicates (data level 3, see below), excluding the comparison of each replicate to itself. Three examples of paired comparisons and the full spectrum of correlations are shown in Figure S1C. We then computed the median correlation between each replicate and all others, yielding 36 values per cell line; and finally summarized using the median of medians so as to derive one value per cell line. These analyses showed that in general the L1000 assay has very high technical reproducibility.

### Reproducibility of L1000—using reference mRNA and cross platform analysis

Samples of purified total RNA from six human cancer cell lines were purchased from Life Technologies. One gene-expression profile per sample was generated using the Affymetrix GeneChip HG-U133 Plus 2.0 Array, the Illumina Human HT-12 v4 Expression BeadChip Array and mRNA-seq (Illumina Hi-Seq) by Expression Analysis, a genomics contract research organization. The L1000 samples were profiled in multiple replicates. Data were normalized within platform (level 3, see below for details). For each cell line, we selected the L1000 replicate with highest technical quality (by LISS goodness of fit, see below) for comparison with the other three platforms. We then performed ComBat batch correction to adjust for cross-platform differences (Johnson, Li, and Rabinovic 2007), and subjected the data to hierarchical clustering in the space of the 952 genes commonly measured by all four platforms. We observe that the data cluster by cell line and not by platform, suggesting that the cross-platform differences are smaller than the biological differences between cell lines.

### Measurement of L1000 using shRNAs

The fidelity of L1000 depends on being able to quantify endogenous levels of intended landmark genes accurately and specifically. In synthesizing landmark gene-specific oligonucleotide probes we followed several computational procedures that maximized matches to the target DNA sequence while minimizing non-specific hybridization. However, as sequence-based QC methods are imperfect and measurement of a transcript might degrade in a multiplexed gene assay (e.g due to cross hybridization), we designed an experiment to empirically confirm probe performance.

To assess the specificity of L1000 landmark probe measurements, we procured shRNAs that target landmark genes from The RNAi Consortium (TRC). We restricted this experiment to shRNAs that had been validated to down-regulate their intended target through RT-PCR assays conducted by TRC. We plated MCF7 and PC3 cells onto 384-well plates and used standard arrayed lentiviral protocols to infect the cells with these shRNAs, each of which targets a specific landmark gene, and then profiled the cells by L1000.

The resulting L1000 signature was used to calculate the targeted landmark gene down-regulation and rank relative to all other shRNAs in the experiment. For each gene in each sample, we computed differential expression values (z-scores) by comparing the gene’s expression value in the given sample to that same gene’s expression values in all other samples in the cohort and then collapsed replicate samples (DS_LM-KD_). The resulting dataset contains, as columns, an individual shRNA targeting a landmark gene performed in either MCF7 or the PC3 cancer cell line. Rows are replicate-collapsed z-scores (level 5, see below) of all measured landmark genes.

A probe designed against a landmark gene was progressed if its z-score when targeted by an shRNA was -2.0 or lower. When the initial probe design showed non-specific reactivity, failed to correlate with reference mRNA standards or failed to register adequate knockdown, we redesigned the probe sequence and retested. After a few cycles of iteration between design and empirical testing, we were able to show that 846 of the 955 targeted landmark genes (89%) were down-regulated by at least one targeting shRNA (z-scores less than -2).

However, a low z-score doesn′t in itself imply specificity—for example, a sample corrupted by dead cells might have yielded low mRNA across the board, leading to many genes with low z-scores. To guard against non-specific reduction of z-scores, we compared the distribution of targeted gene z-scores to non-targeted gene z-scores and observed that the former was significantly left-shifted, indicating that the observed down-regulation is largely specific to the targeted genes (Figure 1C, middle panel). For each targeted gene, we computed the rank of its z-score in the experiment in which it was targeted relative to all other experiments in the dataset where it was not targeted. We observe that 841 of 955 genes (88%) rank in the top 1% and 907 of 955 (95%) rank in the top 5% (Figure 1C, bottom panel). These results indicate that the large majority of L1000 probes are specifically measured.

### Definition of Recall (R)

An absolute measure of similarity (e.g. Spearman correlation) between samples or genes does not in itself convey how uncommon that similarity is. Hence, in addition to computing the similarity (*sim*) between designated samples or genes, it is also useful to compare this similarity value to a reference distribution of similarity values (*SIM*_*null*_), which can aid in interpretation of *sim*. To that end, we compute recall (*R*) as the fraction of *SIM*_*null*_ that is lower than *sim*. High *R* values correspond to unusually high values of *sim*. Thus, *R* provides an assessment of how well a particular pair of samples or genes match each other relative to an appropriate null.

### Reproducibility of L1000—comparisons to RNA-seq from GTEx

We sought to compare expression profiles generated using L1000 with those generated using Affymetrix and RNA-seq, the most widely employed platforms for gene expression profiling. In conjunction with the NIH’s Genotype Tissue Expression (GTEx) project (http://commonfund.nih.gov/GTEx/index), we profiled 3,176 samples on L1000 and obtained from GTEx the RNA-seq (Illumina TrueSeq RNA sequencing) data for different aliquots of these same samples (DS_GEO-RNA-seq_ and DS_GEO-L1000_). The data were quantile normalized independently by platform (level 3, see below) and then batch-corrected using the ComBat algorithm, an empirical Bayes-based method commonly used to remove batch effects across gene expression datasets (Johnson, Li, and Rabinovic 2007).

A small subset of these samples were also profiled on Affymetrix, and Figure 1E, top panel, shows comparisons of the platforms with each other for a single such sample. We observe that L1000 measurements and inferred expression values are as similar with RNA-seq as RNA-seq is with Affymetrix.

To more thoroughly compare L1000 to RNA-seq, we then computed sample self-correlations (using Spearman rank correlation) for the 3,176 samples in the space of the 970 genes directly measured by both platforms. There are 8 L1000 landmark genes that were not included in the DS_GEO-RNA-seq_. Level 3 L1000 data were used, and the GTEx RNA-seq data were quantile normalized, log2 scaled 1+RPKM values. The overlaid histograms (Figure 1E, bottom left) show the distributions of the self-and non-self-correlations (correlations between different samples) for all 3,176 samples. We observe that the 3,176 samples have a median self-correlation of 0.84, and that this distribution is notably right-shifted relative to the non-self-correlations. We then computed the recall (*R*) for each sample and we observed that 3,103 of the 3,176 samples (98%) have a *R* below 0.01, and all but 5 samples (99.84%) have a *R* above 0.95, indicating the the expression profiles generated on L1000 are highly similar with their RNA-seq-derived equivalents (Figure S1D). For each L1000 sample, *R* was computed using the distribution of correlations between the given L1000 sample and all other RNA-seq samples as *SIM*_*null*_.

## 6. EVALUATING GENE INFERENCE

### Gene inference model training

Using the 978 landmark genes as independent variables, we trained an ordinary least squares (OLS) linear regression model on the 12k sample DS_GEO_ dataset, resulting in DS_GEO-OLS_, a matrix of linear coefficients between each landmark gene and each inferred feature. Code is available as above.

The DS_GEO-OLS_ based inference model reports on the 22,268 features (probe sets) measured on the Affymetrix U133A chip, which map to 13,210 unique genes (based on NCBI Entrez Gene as of July 27, 2016). The L1000 assay directly measures 978 probe set ids (corresponding to 978 unique genes) and DS_GEO-OLS_ inference model applied to the L1000 measurements infers the remaining 21,290 probe sets. These 21,290 probe sets correspond to 12,688 unique genes, of which 456 genes overlap with landmark genes—some genes are measured on multiple Affymetrix U133A probe set ids. Taken together, applying the DS_GEO-OLS_ model to a new L1000 sample generates value as below. Gene breakdown of features produced by *infer_ols*:

- Measured: 978
- Measured and inferred (L1000 has a probe set for both): 456
- All inferred (including the 456 that also have a landmark probe set): 12,688
- Inferred only: 12,232
- Measured plus inferred (all genes reported by L1000): 13,210

### Technical note on redundancy in probe sets

To make the output of L1000-based DS_GEO-OLS_ inference more generalizable, we converted it to entrez gene IDs. This was done by comparing each of the inferred probe sets to its measured equivalent, where possible, using a test dataset of GTEx RNA-seq expression profiles (DS_GTEx-RNA-seq_), described in more detail below. This RNA-seq dataset contains measurements for 970 of the 978 landmark genes and 11,350 of the 12,232 inferred genes (corresponding to 17,996 probesets), for a total of 12,320 genes common to both platforms. This cross-platform comparison resulted in a reduction of the feature space of DS_GEO-OLS_ from 22,268 probesets to 12,328 genes -the 12,320 genes common with DS_GTEx-RNA-seq_ as well as the 8 landmarks that were not common.

### Identifying well inferred genes

We sought to assess the inference quality of the 12,232 features corresponding to inferred-only genes in DS_GEO-OLS_. For this test, we used a compendium of 8,555 RNA-seq profiles, generated as part of the GTEx project. We applied the DS_GEO-OLS_ inference model on DS_GTEx-RNA-seq-lmonly_ which resulted in DS_GTEx-RNA-seq-INF_.

To assess inference performance, we computed the correlation of every inferred feature in DS_GTEx-rnase-INF_ to its corresponding gene in DS_GTEx-RNA-seq_. We then analyzed these data to identify genes with statistically significant inferred to measured correlation, as these genes represent the most reliable inference predictions. To generate a null distribution of correlations, we computed the correlation between every inferred probeset in DS_GTEx-RNA-seq-INF_ and every non-matched gene in DS_GTEx-RNA-seq_. We then computed p-values for every inferred gene by computing the percentage of the null distribution with higher correlation than the given inferred gene. We observed that 9,196 of the 11,350 inferred genes (81%) correlated with p-value less than or equal to 0.05. This set of 9,196 inferred genes, plus the 978 landmarks, are referred to as the Best Inferred Genes (BING) and are presented in Table S3.

### Gene space summary

The L1000 assay directly measures 978 genes and infers 11,350 more, for a total of 12,328 genes. Of the 11,350 inferred genes, 9,196 are considered well inferred, based on the analysis described above. All datasets are provided in the full 12,328 gene space. Table S3 indicates which genes are measured or well-inferred.

## 7. GENERATION OF THE FIRST MILLION L1000 PROFILES - EXPERIMENTAL DESIGN

### Overview

L1000 combines locus-specific ligation-mediated amplification with an optically addressed microsphere and flow cytometric detection system to measure selected landmark genes. The result is a 1,000-plex assay at very modest cost which has allowed for the large scale-up reported in this study. However, equally important enablers were process improvements and automation, including:

1. **384-well plates:** Increased sample throughput as opposed to experiments performed on cartridges, glass slides, etc.
2. **Crude cell lysates:** While other plate-based formats for expression screening are available (e.g Affymetrix 96-well plates, HTG plate format), they require extensive upstream sample preparation. In contrast, L1000 input works with whole cell lysate without the need for mRNA purification.
3. **Detection by high speed flow cytometer:** The FlexMAP 3D detection instrument detects 500 bead colors in a high-throughput flow cytometer.

### Control perturbations

The table below describes the control types for compound and genetic experiments. The number of control experiments typically accounted for between 16 and 32 wells on a 384 well plate.

|  | Compound Experiments | Genetic Experiments |
| --- | --- | --- |
| **Positive Controls** | Compounds known to give robust, reproducible signatures (e.g., trichostatin-a) | Reagents targeting landmark genes<br>Reagents targeting well-annotated genes |
| **Negative Controls** | Untreated cells<br>Vehicle treatment only (DMSO) | Untreated cells<br>Vector treatment only<br>Reagents targeting non- |
|  |  | human genes (GFP, RFP, Luciferase, etc.) |

### Replicates

All perturbational profiles were generated in triplicate (minimum of 3 independent treatments of cells with reagent or control). Note that a small number of signatures result from fewer than 3 replicates in cases where one replicate failed due to malfunctioning equipment or other forms of technical error.

### Cell Types

An important goal of the CMap/LINCS undertaking is the collection of data that span different types of perturbations (e.g. genetic and pharmacologic) and that embrace biological complexity and diversity (i.e. are not optimized to one particular model or cell type). In that regard, it would be of value to have a set of perturbations applied to a standard set of cell types (as opposed to different cell lines that are each exposed to a different set of perturbations). The current CMap dataset contains data generated systematically on a set of 9 core cancer cell lines and sparse data on 68 other cell lines representing a mix of cancer, immortalized normal, and primary lines across a diversity of tissue types. To ensure that we are able to generate high-quality L1000 data on a given cell line, we generate baseline data for that cell line and assess the optimal seeding density. We typically test a range of densities and select the one that yields the most robust signal while minimizing dramatic expression changes.

### Selection of compound dose

The selection of appropriate dose is always a topic of intense debate. We believe that there is no single “correct” dose for any given small molecule, and in fact the range of different cellular effects of a compound at different concentrations can be of interest. In time, a full-scale CMap effort might indeed include profiling across a diversity of concentrations. For this initial effort, however, we believed that that community would be best served by profiling a single dose for a large number of pharmacologic perturbations in a large number of cell types (as opposed to a limited number of perturbations studied in detailed dose response). We therefore adopted the following practical strategy. When possible, we used the concentration reported to be effective in cell culture, or the concentration at which the compound scored in a primary screen. Where this information was unavailable, we used a concentration of 10 μM (as is standard practice in most high-throughput small-molecule screens). A subset of our data has been generated at multiple doses and in future profiling efforts will expand the number of doses for many additional compounds.

### Selection of treatment duration

As with dose, there is likely no single “correct” treatment time for compounds or genetic perturbations. We therefore selected a small number of timepoints across which data were generated systematically such that the dataset was more or less standardized along this dimension. For compounds, the treatment times were 6 and/or 24 hours (many compounds were profiled at both timepoints). For genetic reagents, we largely profiled at 96 hours, though a subset of reagents were also profiled at other timepoints to explore their effects.

### Cell culture methods for small-molecule perturbations

Compounds were obtained at 1,000X final concentration in DMSO in 384-well plates from the Broad Institute’s Compound Management Platform and applied to freely cycling human cells also in 384-well plates in accordance with protocols developed for the Connectivity Map.

For treatment, cells were plated into 384-well plates at their optimized seeding densities 6 or 24 hours prior to treatment using a robotic liquid handler. Stock compounds (1,000x) were pinned into the cell culture plate using the CyBio pintool. When treating large batches of plates, we use an intermediate working stock plate (100-fold) diluted in cell culture medium, and then transfer the diluted stocks to the cell culture plates using Cybio 384-well tips (another 10-fold dilution). Treated cells were incubated for 6 or 24 hours, and then lysed by removal of the culture media and addition of TCL lysis buffer using a liquid handling system. Cell lysate plates were sealed using a plate sealer, incubated at room temperature for 30 minutes and then frozen at - 80°C until ready for L1000 profiling.

### Cell culture methods for genetic perturbation experiments

Cell culture and lentiviral transfections for cDNA and shRNA treatments were performed according to the protocols from The RNAi Consortium (TRC), as described in Moffat et al. 2006. CRISPR Cas9 infection and activity assays were performed according to Doench et al. 2014. CRISPR sgRNA cell culture and lentiviral transfections were also performed according to Moffat et al. 2006.

## 8. DATA PRE-PROCESSING - RAW DATA TO SIGNATURES

### Data processing - overview

The L1000 automated data processing pipeline captures raw data from Luminex scanners as it is generated, deconvolutes 978 transcripts from only 500 Luminex bead colors, normalizes the data based on 80 invariant control genes, infers the expression of the non-measured transcripts, determines differentially expressed genes following a perturbation compared to controls, and generates composite signatures across biological replicates. Along the way the data are subjected to rigorous quality control filters at both the sample and plate level.

### Level 1 - Raw (LXB)

Level 1 data comprises the bead identity and raw fluorescent intensity (FI) values measured for every bead detected by the Luminex scanner. The FI is proportional to the amount of amplicon bound to the bead, and hence also proportional to the transcript abundance of the genes that particular bead is interrogating.

### Level 2 - Deconvolute (GEX)

The raw FI values associated with each bead color are analyzed in a peak deconvolution step to associate the expression levels with the appropriate genes. This step is necessary because each bead color is associated with two genes rather than one. To facilitate the analysis, separate bead batches that identify each gene are mixed in a 2:1 ratio for use in the assay. To deconvolute the composite fluorescent intensity signal into its two component expression values and associate them with the appropriate genes, we construct a histogram of FI values. This yields a distribution that generally consists of two peaks, a larger one that designates expression of the gene for which a larger proportion of beads are present, and a smaller peak representing the other gene. Using the k-means clustering algorithm, the distribution is partitioned into two distinct clusters, such that the ratio of cluster membership is as close as possible to 2:1, and the median expression value for each cluster is then assigned as the expression value of the appropriate gene.

### Level 3 - Normalization (NORM)

In order to eliminate artifacts (non-biological sample variation) from the data, we use a rescaling procedure called L1000 Invariant Set Scaling, or LISS, involving 80 control transcripts (8 each at 10 levels of low to high expression) that we empirically found to be invariant in expression across the DS_GEO_. The 80 genes are used to construct a calibration curve for each sample. Each curve is computed using the median expression of the 8 invariant genes at each of the 10 pre-defined invariant levels. We then loess-smoothed the data and fit the following power law function using non-linear least squares regression:

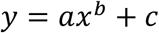

where *x* is the unscaled data and *a*, *b*, and *c* are constants estimated empirically. The entire sample is then rescaled using the obtained model. LISS therefore serves as a method to both adjust for technical variation and to convert between measured Luminex intensity and more traditional Affymetrix log2-expression values.

After applying LISS, we standardize the shape of the expression profile distributions on each plate by applying quantile normalization, or QNORM. This is done by first sorting each profile by expression level, and then normalizing the data by setting the highest-ranking value in each profile to the median of all the highest ranking values, the next highest value to the median of the next highest values, and so on down to the data for the lowest expression level.

Normalization yields the expression values of the 978 landmark genes. To obtain expression values for all the remaining genes in the transcriptome, we assume that an unmeasured gene *x* can be predicted from the measured landmark genes *l*_*i*_ via linear regression:

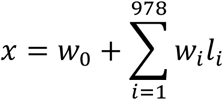

where the *w*_*i*_ constitute the model weights and have been estimated using DS_GEO_. These weights are provided in the dataset *DS*_*GEO-OLS*_. Repeating this procedure for all unmeasured genes gives predicted measurements of all 12,328 genes reported (measured plus inferred) by the L1000 assay.

### Level 4 - Differential Expression (ZSPC)

To obtain a measure of relative gene expression, we use a robust z-scoring procedure to generate differential expression values from normalized profiles. We compute the differential expression of gene *x* in the *i*th sample on the plate as:

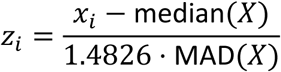

where *X* is the vector of normalized gene expression of gene *x* across all samples on the plate, *MAD* is the median absolute deviation of *X*, and the factor of 1.4826 is a scaling constant to rescale the data as if the standard deviation were used instead of the median absolute deviation.

### Level 5 - Replicate-consensus signatures (MODZ)

L1000 experiments are typically done in 3 or more biological replicates. We derive a consensus replicate signature by applying the moderated z-score (MODZ) procedure as follows. First a pairwise Spearman correlation matrix is computed between the replicate signatures in the space of landmark genes with trivial self-correlations being ignored (set to 0). Then weights for each replicate are computed as the sum of its correlations to the other replicates, normalized such that all weights sum to 1. Finally the consensus signature is given by the linear combination of the replicate signatures with the coefficients set to the weights. This procedure serves to mitigate the effects of uncorrelated or outlier replicates, and can be thought of as a ′de-noised′ representation of the given experiment′s transcriptional consequences.

## 9. CMAP QUERY METHODOLOGY

### Overview of Queries

The fundamental unit of CMap analysis is the query. A query (*q*) consists of a set of genes corresponding to any biological state of interest. Each gene in the query carries a sign indicating if it is up-regulated or down-regulated. Thus each query yields a pair of mutually exclusive gene lists (*q*_*up*_,*q*_*down*_). The query is compared to each signature in the CMap reference database (*Touchstone*) using the similarity metric described below to assess connectivity *viz.* the degree to which the up-regulated query genes (*q*_*up*_) appear toward the top of the rank-ordered signature and the down-regulated query genes (*q*_*down*_) appear toward the bottom of the signature (positive connectivity) or vice-versa (negative connectivity). The result of a query is a rank ordered list of CMap signatures ordered by their connectivity scores.

### Computing similarities - Weighted Connectivity Score (WTCS)

The weighted connectivity score (*WTCS*) represents a non-parametric, similarity measure based on the weighted Kolmogorov-Smirnov enrichment statistic (*ES*) described previously (Subramanian et al. 2005). *WTCS* is a composite, bi-directional version of *ES*. For a given query gene set pair (*q*_*up*_,*q*_*down*_) and a reference signature *r*, *WTCS* is computed as follows:

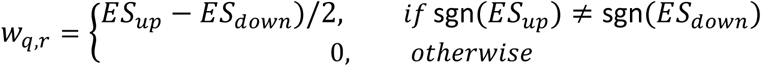

Where *ES*_*up*_ is the enrichment of *q*_*up*_ in *r* and *ES*_*down*_ is the enrichment of *q*_*down*_ in *r. WTCS* ranges between -1 and 1. It will be positive for signatures that are positively related and negative for those that are inversely related, and near zero for signatures that are unrelated. A null (0) score is assigned for cases when both *ES*_*up*_ and *ES*_*down*_ are the same sign.

### Normalization of Connectivity Scores

To allow for comparison of connectivity scores across cell types and perturbation types, the scores are normalized to account for global differences in connectivity that might occur across these covariates. Given a vector of WTCS values w resulting from a query, we normalize the values within each cell line and perturbagen type to obtain normalized connectivity scores (NCS) as follows:

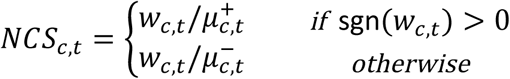

where *NCS*_*c,t*_, *w*_*c,t*_, 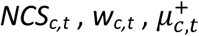 and 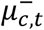 are the normalized connectivity scores, raw weighted connectivity scores, and signed means of the raw weighted connectivity scores (the mean of positive and negative values evaluated separately) within the subset of *Touchstone* signatures corresponding to cell line *c* and perturbagen type *t*, respectively.

Overall, this procedure is similar to that used in Gene Set Enrichment Analysis, with the addition of bidirectional gene sets (i.e up and down) as queries.

### Comparison to reference queries - computation of τ

While meaningful comparisons can be made between the NCS values of reference signatures w.r.t query *q*, it is also useful to assess if the connectivity between *q* and a particular signature *r* is significantly different from that observed between *r* and other queries. This is done by comparing each observed NCS value *ncs*_*q,r*_ between the query *q* and a reference signature *r* to a distribution of NCS values representing the similarities between a reference compendium of queries (*Q*_*ref*_) and *r*. This procedure results in a measure we refer to as τ that ranges [-100, +100] and represents the percentage of queries in *Q*_*ref*_ with a lower |NCS| than |*ncs*_*q,r*_|, adjusted to retain the sign of *ncs*_*q,r*_:

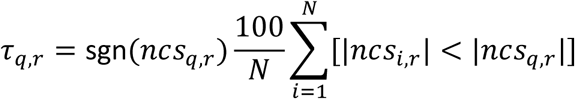

where *ncs*_*q,r*_ is the normalized connectivity score for signature *r* with respect to query *q, ncs*_*i,r*_ is the normalized connectivity score for signature *r* relative to the i-th query in *Q*_*ref*_, and *N* is the number of queries in *Q*_*ref*_ Our standard practice is that *Q*_*ref*_ be comprised of queries obtained from exemplar signatures of *Touchstone* perturbagens that match the cell line and perturbation type of signature *r*. In principle any arbitrary compendium of gene sets (as long as they are large enough) could be used.

### Summarization Across Cell Lines

When examining query results, it is often convenient to obtain a perturbagen-centric measure of connectivity that summarizes the results observed in individual cell types. This can be particularly helpful when searching for connections that persist across cell lines or when one is unsure which cell line to examine. Given a vector of normalized connectivity scores for perturbagen p, relative to query q, across all cell lines in which p was profiled, a cell-summarized connectivity score is obtained using a maximum quantile statistic:

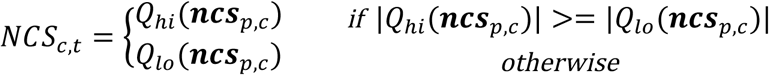

where ***ncs***_*p,c*_ is a vector of normalized connectivity scores for perturbagen p, relative to query q, across all cell lines in which p was profiled, *Q*_*hi*_ and *Q*_*lo*_ are upper and lower quantiles respectively. This procedure compares the *Q*_*hi*_ and *Q*_*lo*_ quantiles of ***ncs***_*p,c*_ and retains whichever is of higher absolute magnitude. Thus, maximum quantile is more sensitive to signal in a subset of the cell lines than measures of central tendency such as mean or median. In the analyses presented here, we used *Q*_*hi*_= *67*, *Q*_*lo*_= *33*

## 10. FEASIBILITY OF QUERYING A MILLION PROFILE COMPENDIUM

When making data at scale, it is important to assess the fidelity of L1000 signatures relative to existing data generated in other labs and on other platforms. This was done by assembling a collection of external (non-L1000) gene sets from GEO and MSigDB and, based on their annotations, computationally determining the list of CMap perturbagens and/or perturbagen sets to which they were expected to connect. Specifically, we scanned the descriptions for 7,578 gene sets and looked for valid CMap perturbagen names. If there was a match, we associated that gene set with the matching given perturbagen (*p*). Gene sets representing compounds were associated with that same compound in CMap and gene sets representing genetic perturbations were associated with either the knockdown and/or overexpression of the same gene in CMap, where possible. Of the 7,578 gene sets, 1,143 matched at least one CMap perturbagen. These 1,143 gene sets correspond to 102 unique compounds and 357 unique genes. One such example is described:

- **Gene set id:** 520CC8E2438AA44B4884A7CB
- **GEO series:** GSE10433
- **Gene set description:** skin, 1 week isotretinoin treatment vs skin, no treatment
- **Associated perturbagen:** isotretinoin

We repeated this procedure systematically and observed that 909 of the 1,143 gene sets (80%) recovered their expected connection at an FDR ≤ 0.25, and |τ| ≥ 90. Full results as well as gene set source information are available in Table S5.

GEO gene sets were derived by comparing the treatment versus control samples within each experiment and selecting the top and bottom 100 BING genes by signal to noise. MSigDB gene sets were simply downloaded as-is and then subset to BING space. Some MSigDB gene sets were two-sided, meaning there was both an up and down set, and some were one-sided, meaning there was just a single gene set with no directionality. In the latter case, the single-sided gene set was queried as the ‘up’ gene set by convention.

p-values were computed relative to a distribution of aggregated NCS values between a collection of 1,000 randomly-generated, two-sided gene sets and *Touchstone*. For each true query (*q*) / TS pert (*p*) combination, we computed the p-value, as the fraction of random queries that had a higher aggregated NCS to *p* than did *q*. We then applied Benjamini-Hochberg p-value adjustment (using the *p.adjust* method in R version 3.2.1) across all true queries *Q* to obtain the FDR. τ was computed relative to the distribution of aggregated NCS values between all true queries *Q* and all *Touchstone* perturbagens. Because the *Q* gene sets were identified computationally and their annotations and definitions were not always consistent (i.e. convention was not always treatment vs. control), and because for many of these gene sets the expected directionality of connection is unclear, we used an absolute τ rather than considering a specific direction.

## 11. DISCOVERING OFF-TARGET EFFECTS OF SHRNAS

The CMap dataset includes signatures for 13,187 shRNAs targeting 3,799 genes across 9 cell lines with multiple hairpins (typically 3) per gene. This large compendium allows us to systematically examine both the intended biological effects of shRNAs as well as their off-target effects. Examination of global distributions of correlations between shRNA signatures revealed that while the correlations between signatures of shRNAs targeting the same gene were higher than a null distribution constructed by sampling random pairs of shRNAs, the correlation between signatures of hairpins that shared the same 6-mer seed sequence were markedly higher. Figure 3A shows data from the A549 cell line. Similar patterns were observed in all cell lines profiled.

In an effort to mitigate the strong off-target effects of shRNAs, we developed an algorithm to produce a Consensus Gene Signature (CGS) that reflects the consistent (and therefore on-target) gene expression effects of shRNAs. To generate a consensus gene signature (CGS), we first create a pairwise Spearman correlation matrix between all shRNA signatures targeting the same gene, explicitly setting self-correlations to 0. Each shRNA signature is then assigned a weight given by the sum of its correlations to the other signatures, with the weights normalized to sum to 1. The CGS is computed as the linear combination of the shRNA signatures, with coefficients set to the weights.

## 12. CHARACTERIZING SMALL-MOLECULE FUNCTION

### Assessing recovery of expected connections

We sought to assess the degree to which each perturbagen profiled in L1000 recovered its expected connections to other perturbagens in *Touchstone* by leveraging annotations compiled from various sources (see 13. Defining and Analyzing Perturbagen Classes). First, the annotations were used to construct a pairwise binary association matrix for all perturbagens in *Touchstone*. A pair of perturbagens were considered to be associated if they shared at least one type of annotation. For example a pair of small-molecules were associated if they shared the same MoA. Similarly a compound and a genetic perturbagen could be associated if they shared the same gene target. We retained 1,902 small-molecule, 994 genetic over-expression, and 1,634 CGS perturbagens after excluding those that had too few (<10) or too many (>3,000) connection pairs. Then for each perturbagen p, we partitioned all associated perturbagen-pairs into a collection of expected connection pairs (*E*_*p*_) whose members were associated with p and and a collection of background pairs (*B*_*p*)_ whose members were not associated with p. Finally, ROC analysis was performed wherein the connectivities between members of *E*_*p*_ were compared to that between members of *B*_*p*_ at different threshold values for connectivity τ ranging (0, 100). At each threshold we computed true positive rates (TPR) as the fraction of *E*_*p*_ that were connected, and false positive rates (FPR) as the fraction of *B*_*p*_ that were connected, thereby generating an ROC curve from which an AUC was derived.

### Target ID Analysis

For this analysis, we evaluated 926 of the 1,902 drugs used in the ROC analysis above. These drugs were each annotated as targeting at least one gene for which we have a CGS in CMap-L1000v1, and collectively they target 283 unique genes. We subjected each drug’s level-5 signature to connectivity analysis and recorded the rank of its target amongst all other CGS in each cell line. We found that 143 of the 283 genes (51%) were amongst the top 100 connections to at least one of their targeting compounds in at least one cell line. Full results are available in Table S7.

We also explored whether failure to recover expected connections might be explained at least in part by cellular context. To address this, we selected 25 compounds lacking strong connections to their expected direct targets in the 9 core cell types, and we re-profiled them in additional 39 cell lines. We then computed the correlation of the compound’s TAS with the gene target’s baseline expression (TAS-GEX correlation) across the 48 total cell lines (9 core plus 39 additional). For each of the 25 compound-target pairs, we re-ranked the target and all genes more connected to the compound by TAS-GEX correlation with the compound. In 12/25 cases (48%), the target gene exhibited a dramatic shift in rank and ranked within or just outside of the top 100 (highlighted in Table S7).

To further explore the known relationship between a compound’s gene expression signature and cell line genotype, we profiled the MDM2 inhibitor AMG-232 in a panel of ten MCF10A isogenic cell lines. We observed AMG-232 had a dramatic reduction in TAS only in the cell line in which *TP53*, which is negatively regulated by *MDM2*, was homozygously deleted compared to the other 9 cell lines, which were all *TP53* wild-type. This result may indicate the utility of a more general screening approach by which the potential target(s) of a compound could be identified by generating L1000 profiles across a diversity of genetic backgrounds.

## 13. DEFINING AND ANALYZING PERTURBAGEN CLASSES (PCLS)

### Overview of PCLs

In order to define perturbational classes we first obtained annotations for as many *Touchstone* perturbagens as possible. For compounds, mechanism of action and gene target annotations were collated from multiple sources (Corsello et al., 2017). For genes, family and pathway annotations were obtained from HGNC as of July 2016. Annotations for both compounds and genes were manually regularized. We next grouped perturbagens by shared annotation to generate candidate classes. For example, all compounds that share the same mechanism of action were assigned to the same class.

For each perturbagen member of a candidate class, we assessed whether it sufficiently recovered its expected connections to other perturbagens in at least one cell line via ROC analysis (more detail below). The class definition was refined to include only those members that passed this criterion. Finally, the classes were assessed for sufficient interconnectivity. We required that classes had at least 3 members and exhibited a median pairwise τ of at least 80 in one or more cell lines. Those classes that passed this filter were codified into perturbagen classes (PCLs). This process resulted in 171 PCLs (92 compound, 60 LoF, and 17 GoF classes) corresponding to 930 unique perturbagens. PCLs ranged in size from 3 to 44 members, with an average size of 5.8 members. PCLs were required to contain only perturbagens of the same type and although perturbagens were allowed to belong to more than one PCL, most PCLs are completely distinct, with a median pairwise overlap of zero members. 95% of PCL members belong to just one PCL.

The majority of PCLs show strong inter-member connectivity in multiple cell types with 132 PCLs (77%) having a cell-summarized median pairwise τ >= 80. 24 PCLs (14%) had significantly stronger connectivity in a particular cell type than in cell-summarized mode, indicating that for these PCLs the connectivity was driven by cell context. Some examples include PPAR receptor agonists in HT29 and PC3 cell lines and estrogen-receptor agonists and antagonists in MCF7.

Compound PCLs were also assessed for structural similarity. The 2D structural similarity of all pairwise combinations of compounds within each PCL was measured using Tanimoto coefficient calculated from binary fingerprints, which were obtained from SMILES strings representing structures of the compounds in PCLs. SMILES strings were converted to binary fingerprints using the Open Babel implementation of the Daylight fingerprint standard (O'Boyle et al. 2011). We found that the vast majority of PCLs were structurally diverse. All but one PCL had a median pairwise Tanimoto below 0.8. Detailed information on all PCLs is available in Supplementary Table S8.

### Computing Connectivity to PCLs

Connectivity of a query to PCLs is computed using the same approach described earlier for summarization of connectivities to a perturbagen across cell lines. Given a vector of normalized connectivity scores for the members of a PCL *p*, relative to query *q*, in a given cell line, we apply the maximum quantile procedure to obtain a summarized *NCS* value (*NCS*_*PCL*_). We then compute a PCL-level τ from *NCS*_*PCL*_ by comparison to a reference distribution comprised of PCL-aggregated scores corresponding to the *Q*_*ref*_ queries described above. This ensures that τ is always computed relative to an equivalent background distribution and keeps it on a scale comparable to that of individual perturbagens.

### Selectivity of PCL Connections

We define the PCL selectivity *s* of a query *q* as the fraction of PCLs whose connectivity to *q* is less than a given threshold *τ*_*th*_. The fewer the number of PCLs connected to by *q*, the higher its selectivity.

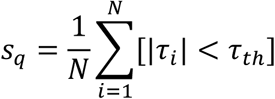

Where *s*_*q*_ is the PCL selectivity of query *q*, *N* is the number of PCLs, τ_*i*_ is the connectivity of *q* to the *i*th PCL and *τ*_*th*_ is the connectivity threshold.

### PCL validation

In order to test the accuracy of PCL connections, we profiled 137 holdout compounds known to share a mechanism with one or more of 54 small-molecule PCLs, but which were not used in the construction of the PCL itself. We subjected the resulting signatures to connectivity analyses as described above and observed that for 41/54 classes (76%), the test compounds connected to their designated PCL in multiple cell types (Figure 4B). For an additional 7/54 (13%), a selective connection was observed in a single cell type. The remaining 6/54 (11%) did not reconnect at a threshold of τ >90. Thus, 48 of the 54 assessed PCLs (89%) were considered validated in that they successfully connected to their corresponding holdout compound(s).

### Unexpected Connections Between Drugs and PCLs

We assessed whether a validated PCL had strong, selective, but unexpected connections to 3,333 annotated small molecule compounds. To focus on unexpected connections, we identified all compounds that connected to a validated PCL at τ ≥ 98 of which the given compound was not a member and whose members’ gene targets did not overlap with the given compounds’ gene targets. For each compound, we computed its PCL selectivity as the fraction of the 171 PCLs to which it failed to connect with τ ≥ 90 and considered only compounds with selectivity of at least 0.9. We identified 225 novel connections between drugs and validated PCLs, corresponding to 132 drugs (3.9% of total assessed). We applied the same analysis to 2,418 unannotated but transcriptionally active compounds and identified 194 strong, selective connections corresponding to 111 compounds (4.6% of total assessed).

### HDAC Inhibitor PCL Clustering

We performed hierarchical clustering on the 22 members of the HDAC inhibitor PCL in the space of their pairwise connectivities to each other across 9 cell lines using spearman correlation as the similarity metric with complete linkage. Hierarchical clustering of pairwise connectivities of the HDAC inhibitor PCL members reveals substructure within the class. The pan-HDAC inhibitors generally cluster together, distinct from the more isoform-selective compounds, suggesting that gene expression can be used to further stratify compounds within the same class (Fass et al. 2010; Bradner et al. 2010; Butler et al. 2010; Kwon et al. 1998; Arts et al. 2009; Haggarty et al. 2003; Bantscheff et al. 2011; Oehme et al. 2013; Balasubramanian et al. 2008)

## 14. CELLULAR CONTEXT

A common question with respect to perturbational signatures is the extent to which they are consistent across different cellular contexts. To investigate this, we first restricted our analysis to the cell lines in which each perturbagen gave a signature whose transcriptional activity score (TAS) was greater than 95% of that of negative controls and considered only perturbagens that had high-TAS signatures in at least three cell lines. Using these thresholds, we analyzed 1,399 of 2,429 compounds, 1,088 of 2,160 cDNAs, and 3,926 of 13,187 shRNAs.

We then computed the pairwise similarity (using *WTCS*) between signatures of the same perturbagen in different cell lines, yielding an *N* x *N* matrix of similarity values, where *N* is the number of cell lines in which the perturbagen gave a high-TAS signature. Next we determined the median WTCS between each cell line and all others, yielding a vector of *N* median *WTCS* values (*WTCS*_*med*_). We then computed the median of medians (*MoM*) and range of *WTCS*_*med*_, yielding *WTCS*_*MoM*_ and *WTCS*_*range*_, metrics which indicate the aggregate similarity and the variability thereof between signatures of the same perturbagen in different cell lines. Perturbagens that give a single signature across multiple cell types should have high *WTCS*_*MoM*_ and low *WTCS*_*range*_ values, respectively. To estimate significance, we computed *WTCS*_*MoM*_ and *WTCS*_*range*_ for 1,000 random combinations of *N* high-TAS signatures for values of *N* between 3 and 9. For a perturbagen to be considered as giving a single signature, we required that its *WTCS*_*MoM*_ be greater than 95% and its *WTCS*_*range*_ be less than 95% of its size-matched null.

Using these thresholds, we found that 26% of compounds, 8% of cDNAs, and 34% of shRNAs gave a single signature across multiple cell lines. The comparatively larger proportion of shRNAs that give a single signature may be attributed to the higher transcriptional activity of shRNAs. We observe that for about 36% of genes with at least 3 high-TAS shRNAs, the majority of those shRNAs were flagged as single-signature reagents. This is not notably different from the 34% rate at which shRNAs give single signatures in general, suggesting that whether or not an shRNA gives a single signature is more dependent on the shRNA itself (and possibly its off-target effects) than it is on the specific gene the shRNA is targeting. cDNAs least frequently gave a common signature. Possibly the high multiplicity of signatures from cDNAs results from a proportionately higher on-target effect which may be more cell context dependent relative to other perturbagen types.

Amongst those perturbagens that were identified as having a single signature, we observed many that target core biological processes such as heat shock response, cell cycle, and HDAC and topoisomerase inhibition, among others. These results suggest that the transcriptional response to perturbing each of these fundamental pathways is conserved across cell contexts.

We also observed a number of classes of perturbagens whose members tended to give multiple unique signatures. For example, 23 of 32 EGFR inhibitors were identified as having multiple signatures and 31 of 34 serotonin receptor antagonists gave multiple signatures, one extreme example being pindolol, whose signature in HCC515 was strongly dissimilar to its signature in other cell lines. These results suggest that the transcriptional response to perturbing these and other pathways may be context-and/or reagent-dependent.

### Neuronal cell line comparison

To extend this analysis to include specialized primary cell types, we considered 768 compounds that had been profiled in both neural progenitor cells (NPC) and differentiated neurons (NEU) as well as the 9 core cancer cell lines. For each compound, we computed the similarity, using WTCS, between all pairwise combinations of cell lines and converted to τ using the pairwise similarities between all 768 compounds in all 11 cell lines as reference (*Qref*). We observed that 189 of the 768 compounds (25%) connected with τ ≥ 90 when comparing NPC to NEU. For each pairwise combination of the 11 cell lines (NPC, NEU + 9 core) we computed the fraction of the 189 compounds that self-connected above 90. We then computed the average fraction that self-connected when considering NPC to cancer (34%), NEU to cancer (25%) and cancer to cancer (50%). This suggests that the neuronal lines are more different from the cancer lines than the cancer lines are from each other, at least in the space of these 189 compounds. Therefore, expanding the cell line set into neuronal cell types may be beneficial.

## 15. IDENTIFYING BIOACTIVE SUBSETS OF SMALL-MOLECULE SCREENING LIBRARIES

The sections and table below describe the various analytical methods used to analyze L1000 signatures. The methods can be broadly grouped into two categories - characterization and connectivity. The characterization methods enable nuanced analyses of L1000 signatures in and of themselves, while the connectivity methods provide a framework by which external gene sets or L1000 signatures can be queried against *Touchstone*, and by which an analyst can derive connectivities to individual perturbagens and perturbagen classes (PCLs).

### Replicate Correlation (CC)

Each L1000 experiment consists of multiple biological replicates. To derive an aggregate measure of replicate reproducibility, we compute the 75th quantile of the Spearman correlations between all pairwise combinations of replicate level 4 profiles for a given experiment.

### Signature Strength (SS)

We compute signature strength (*SS*) as the number of differentially expressed genes within a signature. That is, the number of landmark genes with absolute z-score greater than or equal to 2. The z-scores are adjusted to offset shrinkage of z-scores that occurs with increasing number of replicates. This allows *SS* values derived from signatures of different numbers of replicates to be compared with each other

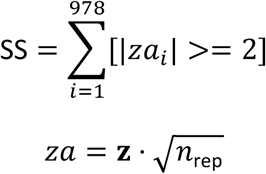

Where **z**, n_rep_ are a vector of moderated z-scores and the number of replicates, respectively.

### Transcriptional Activity Score (TAS)

The transcriptional activity score (*TAS*) is computed as the geometric mean of *SS* and *CC* for a signature. *TAS* is scaled by the square root of the number of landmark genes (978) so the final score ranges between 0 and 1.

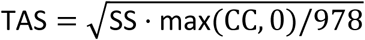

Where *SS* and *CC* are the signature strength and replicate correlation for the given signature, respectively.

### Analysis of unannotated small-molecule screening libraries

We began with a collection of 16,527 unannotated small molecules for which we had generated L1000 profiles. These compounds were derived from a variety of sources, including the Broad Institute’s diversity oriented synthesis (DOS) library and the NIH′s Molecular Libraries Probe Production Centers Network (MLPCN). We focused on the 2,418 compounds whose 75th quantile of TAS was as least 0.2 and whose signatures had a median pairwise WTCS of at least 0.3 across cell lines, indicating a robust transcriptional response in at least a subset of cell lines. We termed these compounds *Discovery*, and attempted to assign functional annotations via comparison with annotated drugs and genes in the L1000 *Touchstone* (reference) part of the data.

To obtain a high-level view of these *Discovery* compounds relative to known drugs, we ran t-SNE analysis on the *Discovery* signatures and those of every compound belonging to a PCL. t-SNE is a non-linear dimensionality reduction and visualization technique that attempts to preserve local-structure from high-dimensional datasets ensuring that samples that are similar in the high dimensional space are plotted close together in the embedding (van der Maaten 2008). t-SNE was run on consensus signatures across cell types for each perturbagen in landmark space, with initial dimensions set to 50 and a perplexity of 30.

In addition, we performed query analysis on these compounds’ signatures to derive their connectivities to *Touchstone* perturbagens and PCLs. We found that 111 *Discovery* compounds had strong and selective connections to PCLs (τ >= 98; PCL specificity >= 0.9).

### Discovery of a selective CSNK1A1 inhibitor

We observed that the unannotated *Discovery* compound BRD-1868 connected to the shRNA-based CGS of *CSNK1A1* with τ ≥ 90 in both A375 and HA1E. We separately generated CRISPR signatures of *CSNK1A1* knockout and observed that BRD-1868 ranked in the top 2% by WTCS similarity to the *CSNK1A1* knockout signature relative to all Discover compounds in two additional cell lines (MCF7 and HT29).

## 16. USING L1000 TO ASSESS ALLELE FUNCTION

A series of variant alleles of genes frequently mutated in primary lung adenocarcinomas were overexpressed in human cell lines as detailed in (Berger et al. 2016) and subjected to L1000 profiling. Connectivites to perturbagens in *Touchstone* were computed using queries derived for each allele. The differences or similarities in connections to specific perturbagens between wildtype and mutant alleles were examined.

## 17. USING CMAP TO INTERPRET CLINICAL TRIAL RESULTS

In the first studies, 21 patients with melanoma were treated with the RAF inhibitor dabrafenib or vemurafenib and 9 patients were treated with dabrafenib plus the MEK inhibitor trametinib (Carlino et al. 2013; Long et al. 2014). Biopsies were obtained prior to treatment and at the time of relapse, and in four patients, early on-treatment biopsies were also obtained. The authors performed expression profiling on the Illumina beadchip platform, and we used those data (GSE50509, GSE61992). In order to evaluate the transcriptional changes induced by *in vivo* drug treatment, we focused on the four patients across the two datasets who had matched pre-treatment, early on-treatment, and relapse biopsies performed. Gene expression results were obtained as series matrix files from GEO and the log2 fold change in each gene was calculated for each sequential comparison (early on-treatment versus pre-treatment and relapse versus early on-treatment). Genes were mapped to CMap best inferred gene (BING) space and the top 100 genes up-and down-regulated in each comparison were used to perform CMap queries. Results are shown for connections within the A375 melanoma cell line given contextual relevance.

In the second study, patients with solid tumors were treated with the pan-CDK inhibitor PHA-793887 in a phase I clinical trial. Seven patients from that trial were subjected to gene expression profiling of biopsies obtained pre-treatment and on-treatment using an Agilent microarray platform (Massard et al. 2011; Locatelli et al. 2010). Gene expression data was obtained as a GEO series matrix (GSE18553). The log2 fold change in each gene was calculated between the early on-treatment and pre-treatment biopsies. Genes were mapped to CMap L1000 landmark gene space and the top 50 genes up-and down-regulated in each comparison were used to perform CMap queries. Connectivity results were reviewed at the cell-summarized level to identify consistent connections across multiple cell types. There was no obvious correlation between medication dose administered or primary tumor type with the *CDK4* shRNA connection, although the number of samples was small.

